# Beyond DNA Binding: single C2H2 zinc fingers with adjacent β-strands mediate dimerization in *Drosophila* transcription factors

**DOI:** 10.1101/2025.09.04.674178

**Authors:** Konstantin I. Balagurov, Sofia S. Mariasina, Sergey D. Dukhalin, Nikolai N. Sluchanko, Anastasia M. Khrustaleva, Alexandra A. Golovnina, Oksana G. Maksimenko, Olga V. Arkova, Alexandra A. Stepanenko, Eduard V. Bocharov, Vladimir I. Polshakov, Pavel G. Georgiev, Artem N. Bonchuk

**Author notes:** To whom correspondence should be addressed (K.I.B.), (A.N.B.).

## Abstract

C2H2 proteins, characterized by DNA-binding C2H2-type zinc finger domains, constitute the largest group of transcription factors. In addition to binding DNA, C2H2 domains can mediate protein–protein interactions, facilitating oligomerization of C2H2 proteins. In this study, we identified eight C2H2 proteins in the *Drosophila* genome that feature a unique single C2H2 domain containing a conserved’CGEx’ motif. Yeast two-hybrid assays, SEC-MALS, and chemical cross-linking experiments revealed a strong propensity of these domains to form dimers. Using NMR spectroscopy, we determined the solution structure of the dimeric C2H2 domain from IMZF (Immune-mediated Zinc Finger) protein, providing structural evidence towards the dimerization of C2H2 domains. The structure revealed that dimerization is mediated by the interface between the core C2H2 fold and the adjacent β-strand containing’CGEx’ motif, which was further validated by structure-guided mutagenesis. A bioinformatic survey showed that’CGEx’-type C2H2 domains are specific to *Diptera*. Finally, we demonstrate that putatively dimerizing C2H2 domains containing additional structural elements beyond the canonical fold are widespread among diverse eukaryotic taxa, with the highest prevalence observed in insects. These findings establish that single C2H2 domains can mediate self-association and identify the’CGEx’-type C2H2 domain as a distinct structural subclass specific to Dipteran insects.

## INTRODUCTION

In higher eukaryotes, proteins with C2H2-type zinc-finger domains (C2H2 proteins) constitute a large group of transcription factors that play a critical role in the regulation of gene expression (1–3). C2H2 domains were first described as sequence-specific, zinc-coordinating DNA-binding modules in the TFIIIA transcription factor (4). Since then, numerous zinc finger subclasses have been identified, distinguished by additional secondary structure elements or variations in zinc-coordinating amino acid residues (2, 5–7). A classic (TFIIIA-type) C2H2 domain consists of two antiparallel β-strands followed by an α-helix, stabilized by two cysteine and two histidine residues coordinating a zinc ion (4, 5). DNA-binding activity of C2H2 domains is primarily determined by the amino acid residues in the α-helix region (8–11).

C2H2 proteins underwent rapid expansion over the course of evolution and are mostly prevalent in higher eukaryotes. For instance, the genomes of *H. sapiens* and *D. melanogaster* contain over 1,700 and 400 proteins of this group, respectively (2). These proteins commonly feature two or more DNA-binding C2H2 domains arranged in clusters, which facilitate the recognition of long, specific binding sites (1, 2, 12–14). In addition to DNA-binding C2H2 domains, some C2H2 proteins possess an N-terminal protein–protein interaction effector domain like BTB (Bric-a-brac, Tramtrack and Broad Complex) (15–20), ZAD (zinc finger-associated domain) (21, 22), SCAN (SRE-ZBP, CTfin51, AW-1, and Number 18 cDNA) (23), and KRAB (Krüppel-associated box) (24, 25). These effector domains mediate the recruitment of interaction partners, promote oligomerization, modulate DNA-binding specificity, and regulate transcription factor activity (26–29). Notably, many C2H2 proteins are devoid of dedicated effector domains, instead many of them feature the presence of isolated C2H2 domains outside of C2H2 clusters (2). The question arises whether these domains might function as effector domains.

Functions of the C2H2 domains are not limited to DNA binding; several studies have demonstrated their ability to mediate various protein–protein interactions (30, 31). Some C2H2 domains form complexes with structural motifs like coiled-coil (32) or engage in intramolecular interactions within the same polypeptide chain (33, 34). Ubiquitin-binding zinc fingers (UBZ), which are related to C2H2 domains, bind directly to ubiquitin (35, 36). Besides this, double C-terminal C2H2 domains in the Ikaros family transcription factors mediate selective homo-and heterodimerization as well as the formation of higher-order complexes (37–42). The self-association of these double C2H2 domains was reported to be important for high-affinity DNA binding and overall functional activity of these proteins (43). However, the structural mechanisms underlying C2H2 domains self-association remain largely unexplored. Structural data on the direct interaction between two different zinc finger domains are limited to atypical zinc fingers, such as C4 zinc finger of GATA-1 interacting with C3H zinc finger of FOG protein (44). Given the available data on C2H2 domain mediated protein– protein interactions and their abundance in higher eukaryotic genomes, it is likely that homo-and heterotypic interactions between C2H2 domains could be widespread, though remaining almost uncharacterized yet (2, 30, 31, 44).

In this study, we screened the *Drosophila melanogaster* proteome for C2H2 proteins that contain a single N-terminal C2H2 zinc finger as a potential self-associating domain, alongside with a cluster of C2H2 zinc fingers. We identified a family of *D. melanogaster* C2H2 transcription factor proteins with an N-terminal dimerization C2H2 domain, comprising eight members: CG1602, Mzfp1/CG1603, CG1605, CG2129, CG10959, IMZF/CG18262, CG8643, and CG8944. The functions of two members of this family (Mzfp1 and IMZF) have been described recently (45–47).

Using yeast two-hybrid assay, glutaraldehyde crosslinking, and SEC-MALS, we demonstrated that all these C2H2 domains exist in a dimeric state in solution. According to phylogenetic analysis, they are found exclusively within the proteins of Dipteran insects. Finally, we determined the molecular structure of the N-terminal C2H2 domain of the IMZF protein using NMR, providing the structural data for the dimerization interface between single C2H2 domains.

## MATERIALS AND METHODS

### Plasmids and cloning

cDNAs for all protein fragments were PCR-amplified with corresponding primers and cloned into modified pET32a(+) vector (Novagen) containing the TEV protease cleavage site after thioredoxin. DNA constructs for expression of mutant proteins were prepared with PCR-directed mutagenesis using corresponding mutagenic primers. For yeast two-hybrid assays, fragments were amplified using corresponding primers and cloned into pGAD424 and pGBT9 vectors (Clontech), containing the activation or the DNA-binding domain of GAL4, respectively. Additionally, fragments were cloned into a modified pGBT9 (pGBT-C) vector containing a multiple cloning site upstream of the GAL4 DNA-binding domain. The sequences of all primers used are listed in **Supplementary Table S1.**

### Protein expression and purification

BL21 (DE3) cells were transformed with modified Pet32a vectors encoding cDNAs of IMZF^1-62^, CG10959^1-75^ or 1605^1-62^ fused to TEV-protease cleavable 6xHis-tag and thioredoxin tag. Cells were grown in 500 ml of LB medium at 37°C until reaching OD=1.0, then ZnCl_2_ was added to a final concentration of 0.1 mM, and expression was induced with 0.8 mM IPTG. Proteins were expressed for 16 hours at 18°C. Mutant forms of IMZF^1-62^ were expressed under the same conditions. Isotopically labeled IMZF^1-62^ for NMR experiments was expressed using an optimized protocol from Marley et al. (48) and purified with the same procedure as unlabeled proteins.

Cells were disrupted using high-pressure homogenizer (Microfluidics) in 15 ml lysis-buffer (30 mM HEPES-NaOH pH 7.5, 500 mM NaCl, 10 mM Imidazole, 0.1 % NP40, 5% (v/v) Glycerol, 0.1 mM ZnCl2, 1 mM DTT, 1 mM PMSF and Calbiochem Complete Protease Inhibitor Cocktail VII (1μL/1ml)). Each lysate was centrifuged at 25000g for 1 hour then applied to a gravity-flow Ni-NTA column, washed with 10 volumes of wash buffer (30 mM HEPES-NaOH pH 7.5, 500 mM NaCl, 30 mM Imidazole, 5% (v/v) Glycerol, 0.1 mM ZnCl2, 1 mM DTT) and eluted with 10 ml of elution buffer (30 mM HEPES-NaOH pH 7.5, 500 mM NaCl, 5% (v/v) Glycerol, 300 mM Imidazole, 1 mM DTT). To cleave 6xHis-thioredoxin tag, approximately 150 μg of 6x-His-tagged TEV protease was added to the eluted protein mixture, and sodium citrate was added to a final concentration of 5 mM. The mixture was dialyzed overnight at 4°C against degassed 30 mM HEPES-NaOH pH 7.5, 500 mM NaCl, 10 mM Imidazole, 5% (v/v) Glycerol, 0.1 mM ZnCl_2_, 1 mM DTT, then filtered and again applied to the gravity-flow Ni-NTA column. Flow-through, containing the cleaved protein, was collected and dialyzed overnight at 4°C against degassed 20 mM Tris-HCl pH 7.4, 50 mM NaCl, 0.1 mM ZnCl2, 1 mM DTT. The protein sample was next purified using MonoQ 5/50 GL (Cytiva). IMZF^1-62^, its mutant forms, and CG10959^1-75^ were collected as flow-through, CG1605^1-62^ was eluted with 0-500 mM gradient of NaCl. In case of CG1605^1-62^, samples with protein of interest were combined, and ammonium sulfate was added to final concentration of 1.5M. Then protein mixture was applied to HiTrap Phenyl HP column (Cytiva), and protein was eluted with 20 mM Tris-HCl pH 7.4, 0.1 mM ZnCl_2_, 1 mM DTT.

For SEC-MALS experiments, purified proteins were dialyzed overnight at 4°C against degassed 20 mM Tris-HCl pH 7.4, 150 mM NaCl, 0.1 mM ZnCl2, 1 mM DTT. For NMR experiments, purified IMZF^1-62^ was dialyzed overnight at 4°C against degassed 20 mM sodium phosphate pH 7, 50 mM NaCl, 5 mM DTT.

### NMR-solution structure of IMZF^1-62^

NMR samples were prepared at concentrations of 0.73 mM for uniformly ^13^C,^15^N-labeled IMZF^1-62^ and 1.0 mM for ^15^N-labeled IMZF^1-62^, in 20 mM sodium phosphate buffer (pH 7.0) containing 50 mM NaCl and 0.02% (w/v) sodium azide. Samples were prepared either in 95% H₂O/5% D₂O or 100% D₂O (for exchange experiments).

NMR spectra were recorded at 25 °C on Bruker AVANCE 600 MHz spectrometers equipped with either a TXI triple-resonance probe (^1^H, ^13^C, ^15^N) or a quadruple-resonance cryoprobe (^1^H, ^13^C, ^15^N, ^31^P). All 2D and 3D spectra were processed using NMRPipe (49) and analysed with POKY (50).

Backbone resonance assignments of ^1^H, ^13^C, and ^15^N nuclei were performed using standard triple-resonance experiments: HNCO, HN(CA)CO, HNCOCA, HNCA, and C(CO)NH. Manual assignment was carried out using POKY, supported by automated predictions from the PINE algorithm (51). Backbone amide ¹H and ^15^N chemical shifts were assigned for all non-proline residues except V4 and Q5 (absent from the spectra), and S58, S60, and S61 (excluded due to peak overlap).

Side-chain assignments were performed manually in POKY using the following experiments: HCCH-TOCSY, DQF-COSY, HNHA, and HNHB. Stereospecific assignments were obtained using a combination of HNHA, HNHB, ^15^N–^1^H HSQC-TOCSY, ^13^C–^1^H HSQC-NOESY, and ^15^N–^1^H HSQC-NOESY spectra, with assistance from the AngleSearch program (52).

### NMR restraints collection and structure calculation

The solution structure of the IMZF^1-62^ homodimer was calculated using the ¹H– ¹H distance restraints derived from NOE data, torsion angle restraints derived from ³J coupling constants and TALOS-N (53) secondary structure predictions, hydrogen bond restraints inferred from hydrogen–deuterium (H/D) exchange experiments, Zn-coordination restraints, and chemical shifts restraints.

### (1) ^1^H-^1^H Distance restraints

Proton–proton distance restraints were derived from the following NMR experiments: 3D ^13^C–^1^H HSQC-NOESY, 3D ^15^N–^1^H HSQC-NOESY and 2D ¹H–¹H NOESY recorded in either 100% D₂O or 95% H₂O/5% D₂O. All NOESY spectra were acquired with a mixing time of 100 ms, resulting in a total of 1674 NOE-derived distance restraints. The number and distribution of restraints are given in **Supplementary Table S2A** and illustrated in **Supplementary Figure S1**. Spectral assignment and structure calculation were carried out iteratively using the ARIA 2.3 software package (54) followed by manual verification and refinement.

### Intermolecular NOE restraints

were specifically obtained from 2D and 3D ^13^C/^15^N-filtered NOESY spectra recorded with a mixing time of 100 ms on a 1:2 molar mixture of uniformly ^13^C/^15^N-labeled and unlabeled IMZF^1-62^. This isotopic filtering approach selectively detects intermolecular NOEs between the labeled and unlabeled chains while suppressing intramolecular NOEs. The 1:2 ratio was used to enhance detection of intermolecular contacts and reduce spin diffusion artifacts.

### (2) Torsion angle restraints

Torsion angle restraints were derived from ³J coupling constants measured in HNHA and DQF-COSY spectra and NOEs measured in NOESY spectra, using the AngleSearch program (52). An additional set of backbone φ and ψ dihedral angle restraints was determined from the chemical shift values of the backbone atoms ^13^Cα, ^13^Cβ, ^13^CO, ^1^Hα, ^1^HN, and ^15^N using TALOS-N software (53).

### (3) Hydrogen bonds restraints

Hydrogen bond restraints were derived from H/D exchange experiments. A freeze-dried ^13^C, ^15^N-labeled IMZF^1-62^ sample (1.0 mM) was re-dissolved in 100% D₂O, and ^1^H–^15^N SOFAST-HMQC spectra were recorded after 1 and 24 hours. The spectra were compared with a reference spectrum acquired in 95% H₂O/5% D₂O. Slowly exchanging amide protons — those showing retained intensity after 24 hours — were identified as candidates for hydrogen bonding. Hydrogen bond restraints were assigned to residues with slow H/D exchange and located near acceptor carbonyl groups in preliminary structural models.

Inter-subunit hydrogen bond restraints were identified from the NMR-derived dimer model based on consistent H/D protection and proximity of potential donor/acceptor pairs across the interface. A total of 6 inter-subunit hydrogen bonds were included in the final restraint set.

### (4) Zn-coordination restraints

Zn-coordination restraints were applied to maintain proper geometry of the zinc-binding sites in IMZF^1-62^. The tetrahedral coordination of Zn^2+^ ions by conserved Cys and His residues was enforced using distance restraints between the metal center and coordinating atoms (Cys S_γ_ and His N_ɛ_). These restraints were derived from QM/MM calculations on similar zinc finger domains (55) and validated by the absence of NOE violations upon Zn^2+^ chelation.

### (5) Chemical shifts restraints

Chemical shift restraints for ^1^Hα atoms were incorporated into the structure calculations as harmonic pseudo-energy terms during simulated annealing refinement using the CNS program.

Final restraint summary is shown in the **Supplementary Table S2A**.

### Structure Calculation and Refinement

Structure calculations were performed using a simulated annealing protocol in Cartesian coordinate space implemented in the CNS 1.21 software package (56). During the final stages of the refinement, database-derived conformational torsion angle pseudopotentials were applied to improve the accuracy of backbone geometry (57).

The overall quality of the calculated structures and restraint violations were evaluated using CNS validation tools, PROCHECK-NMR (58), and the NMRest in-house program (59). Out of 200 calculated conformers, the 20 lowest-energy structures, which also showed no residues in disallowed regions of the Ramachandran plot, were selected to represent the final NMR ensemble of IMZF^1-62^. Structural statistics is shown at **Supplementary Table S2B**.

All non-glycine residues in the final ensemble fall within favored regions of the Ramachandran plot (**Supplementary Figure S2**), indicating high stereochemical quality of the model. A comprehensive summary of the applied restraints and structural statistics is provided in **Supplementary Table S2B.**

### Structure visualization and analysis

were carried out using PyMOL 2.3.0 Open-Source (Schrödinger, LLC) and UCSF ChimeraX 1.8 (Resource for Biocomputing, Visualization, and Informatics, University of California, San Francisco) (60).

### NMR Relaxation, Backbone Dynamics and Rotational Diffusion Analysis

Backbone dynamics were characterized by measuring longitudinal (R₁) and transverse (R₂) relaxation rates using 2D [^1^H–^15^N] correlation spectra acquired with sensitivity-enhanced pulse sequences (61).

For R_1_ measurements, spectra were recorded with relaxation delays of 100, 200, 400, 600, 800, 1000, 1200, and 1500 ms. R_2_ values were determined using CPMG-based spin-echo experiments with delays of 0.16960, 0.01696, 0.13568, 0.05088, 0.11872, 0.06784, 0.08480, and 0.10176 ms. Steady-state {^1^H–^15^N} heteronuclear NOE values were obtained from spectra recorded with and without 3 s of proton saturation, with a 5 s recycle delay between scans. All relaxation experiments were performed at 298 K on a 600 MHz Bruker spectrometer using uniformly ^15^N-labeled IMZF^1-62^ protein samples.

Spectra were processed using NMRPipe (49), and preliminary estimates of model-free parameters (S^2^, Rₑₓ) were obtained using in-house written RelaxFit program (62). Final analysis, including rotational diffusion tensor calculation and refinement of internal dynamics, was carried out using TENSOR2 (63).

The rotational diffusion tensor was computed using residues from the rigid core of the protein (S² > 0.8). An anisotropic diffusion model provided significantly better agreement with experimental data (χ^2^ = 376) than the isotropic model (χ^2^ = 2112), consistent with the non-spherical shape of the dimeric IMZF^1-62^. The final diffusion tensor parameters (D_x_ = 1.63 × 10^7^ s^-1^, D_y_ = 1.85 × 10^7^ s^-1^, D_z_ = 3.20 × 10^7^ s^-1^) revealed pronounced anisotropy (D_ǁ_/D_⊥_ ≈ 2.0), with the principal diffusion axis aligned along the dimer interface.

Model-free analysis of internal motions was refined in TENSOR2. For each residue, five dynamic models were tested, and model selection was based on F-tests (p < 0.2, TENSOR2 default) and Monte Carlo error analysis (100 iterations). Residues showing significant contributions from chemical exchange were identified by comparing models with and without the R_ex_ term; residues with R_ex_ > 1.5 s^-1^ were considered indicative of slow conformational exchange.

### Bioinformatics

A primary search for orthologs of the studied *D. melanogaster* C2H2 proteins was performed in Metazoa using the OrthoDB (64) and EGGNOG databases (65). The total number of identified orthologs from these two combined databases was 1186 (781 from OrthoDB and 405 from EGGNOG respectively). To retrieve IDs from the NCBI database for each orthologous protein, E-Utilities’ EDirect tools, esearch and efetch, were used (66). If the corresponding IDs were not found in the NCBI database, the original identifier from the OrthoDB or EGGNOG database was retained. Since all studied proteins are orthologs for each other, the dataset contained duplicates that were removed with SeqKit2 tool (67). After removing duplicates, the total number of orthologs was reduced to 630. The amino acid sequences of orthologous proteins corresponding to 8 studied *D. melanogaster* C2H2 proteins were aligned using MUSCLE (68). Orthologous proteins that lacked the N-terminal ‘CGEx’ motif were excluded from further analysis. The initial MSA file was trimmed in Jalview (69) to retain only the ‘CGEx’ motif and the subsequent core part of the C2H2 domain. This region was then searched in the UniProtKB database using HMMSEARCH (web-version) (70). 324 sequences of orthologous proteins from the HMMSEARCH output were aligned using MUSCLE and subsequently trimmed in Jalview following the same approach as the initial MSA. AlphaPulldown (71) predictions were then performed for each protein dimer to assess its dimerization potential (**Supplementary File 1**). The final dataset consisted of 449 sequences from 55 Diptera species (**Supplementary File 2**). The resulting multiple sequence alignments of full-length C2H2 orthologs were trimmed with trimAl to remove poorly aligned regions (72).

The taxonomic hierarchy was reconstructed according to the NCBI Taxonomy Database using Taxize (73). Phylogenetic relationships between the proteins were inferred using maximum likelihood (ML) tree reconstruction in IQ-TREE 2.0.3 (74) using General ‘Variable Time’ matrix (75) with discrete Gamma (76) (VT+G4) as a best-fit model by all criteria (AIC, AICc, BIC) calculated in ModelFinder module implemented in IQ-TREE (**Supplementary File 2**).

Relative gene expression analysis was performed using RNA-seq datasets from the modENCODE (RPKM values) (**Supplementary File 2**) (77). Levels of the gene expression were quantified across all major development stages of *D. melanogaster*, as defined by FlyBase (78).

AlphaFold predictions were run locally using AlphaFold 2.3 installation with a full set of genetic databases (79). Large-scale predictions of multimeric status of C2H2 domains with additional secondary structure elements were run using AlphaPulldown pipeline (71).

### Glutaraldehyde cross-linking assays

Vectors encoding cDNAs of all investigated C2H2 domains (Mzfp1^1-47^, CG1602^1-56^, 1605^1-62^, IMZF^1-62^, CG10959^1-75^, CG2129^1-56^, CG8643^1-62^, CG8944^1-80^) fused to TEV-protease cleavable 6xHis-tag and thioredoxin were expressed and purified using a batch method with the same procedure described above, excluding TEV-cleavage and subsequent steps. The empty vector expressing 6xHis-tag and thioredoxin alone was used as negative control. Concentration of protein samples was adjusted to 100 μg/ml. The reactions were performed in a buffer containing 30 mM HEPES-NaOH pH 7.5, 500 mM NaCl, 300 mM Imidazole, 1 mM DTT, and either 0.1% or 0.01% (w/v) glutaraldehyde, or no crosslinker as control. After 10 minutes of incubation, the reactions were terminated by adding Tris-HCl pH 6.8 to a final concentration of 50 mM. Samples were analysed by SDS-PAGE and visualized by silver staining.

### Yeast two-hybrid assays

Yeast two-hybrid assays were performed using optimized protocols from Clontech. Plasmids were co-transformed into the yeast strain pJ69-4A using the lithium acetate method. Yeast cells were grown overnight at 30°C shaking at 250 rpm in 5 mL YPD medium supplemented with 0.003% (w/v) adenine. The overnight culture was diluted 1:10 in fresh medium and incubated for an additional 2 hours under the same conditions. Cells were harvested by centrifugation at 500g for 5 minutes, resuspended in 20 mL of 0.1 M lithium acetate, and incubated with shaking at 30°C for 20 minutes. Then the cells were centrifuged again under the same conditions and resuspended in 15-30 mL of 0.1 M lithium acetate solution containing 50% (w/v) PEG 4000. Approximately 1 μg of each plasmid DNA was added to a 1.5 mL tube, followed by the addition of 100-300 μL of the yeast cells. The cells and the plasmid DNA were incubated at 30°C for 30 minutes with shaking at 250 rpm, heat-shocked for 15 min at 42°C, chilled on ice for 2 minutes, centrifuged at 4000g for 30 seconds, resuspended in 35 μL of sterile water, and plated onto SD media lacking tryptophan and leucine. After 3 days of growth at 30°C, colonies were plated on selective media without tryptophan, leucine and histidine. The cell growth was compared after 3 days. Each assay was repeated three times.

## SEC-MALS

In order to determine the oligomerization state of the zinc fingers in solution, we measured their absolute molar mass by multi-angle static light scattering coupled with size-exclusion chromatography (SEC-MALS), this does not depend on a protein shape or possible interactions of the proteins with the chromatographic resin. Protein samples were first centrifuged for 5 min at 21 400g at 4°C and then were loaded using a Vialsampler (G7129A, Agilent) onto a Superdex 75 10/300 column (GE Healthcare) pre-equilibrated with SEC buffer (20 mM Tris–HCl, pH 7.4, 200 mM NaCl, 0.1 mM ZnCl_2_ and 5 mM DTT). The column was operated at a 1.0 mL/min flow rate using an Agilent 1260 Infinity II chromatography system equipped with a 1260 Infinity II WR diode-array detector (G7115A, Agilent), a miniDAWN detector (Wyatt Technology), and a refractometric detector (G7162A, Agilent, operated at 35 °C) connected sequentially in the indicated order. The elution profiles were followed by absorbance at 280 nm and by changes of the refractive index and the static light scattering intensity at three angles (45°, 90° and 135°) at ambient temperature. The SEC-MALS data were analysed in Astra 8.0 software (Wyatt Technology) using the refractometric detector as a concentration source (dn/dc was taken equal to 0.186 for each protein). Normalization of the light scattering intensities measured at different angles was performed using a pre-run profile of a BSA standard (Wyatt).

## RESULTS

### Single N-terminal C2H2 domain defines the new family of *Drosophila melanogaster* C2H2 transcription factors

C2H2 proteins typically feature clusters of DNA-binding C2H2 domains, often accompanied by a protein–protein interaction domain, commonly mediating oligomerization and located at the N-terminus (2, 25). Thereby, single N-terminal C2H2 domains outside of clusters could have potential self-association properties. To identify such C2H2 proteins, we screened the InterPro database for *D. melanogaster* proteins containing both clustered C-terminal C2H2 domains and a single N-terminal C2H2 domain within a 1-100 amino acid range (80). Our survey identified 8 proteins with such domain organization (**Figure 1A**).

**Figure 1.**
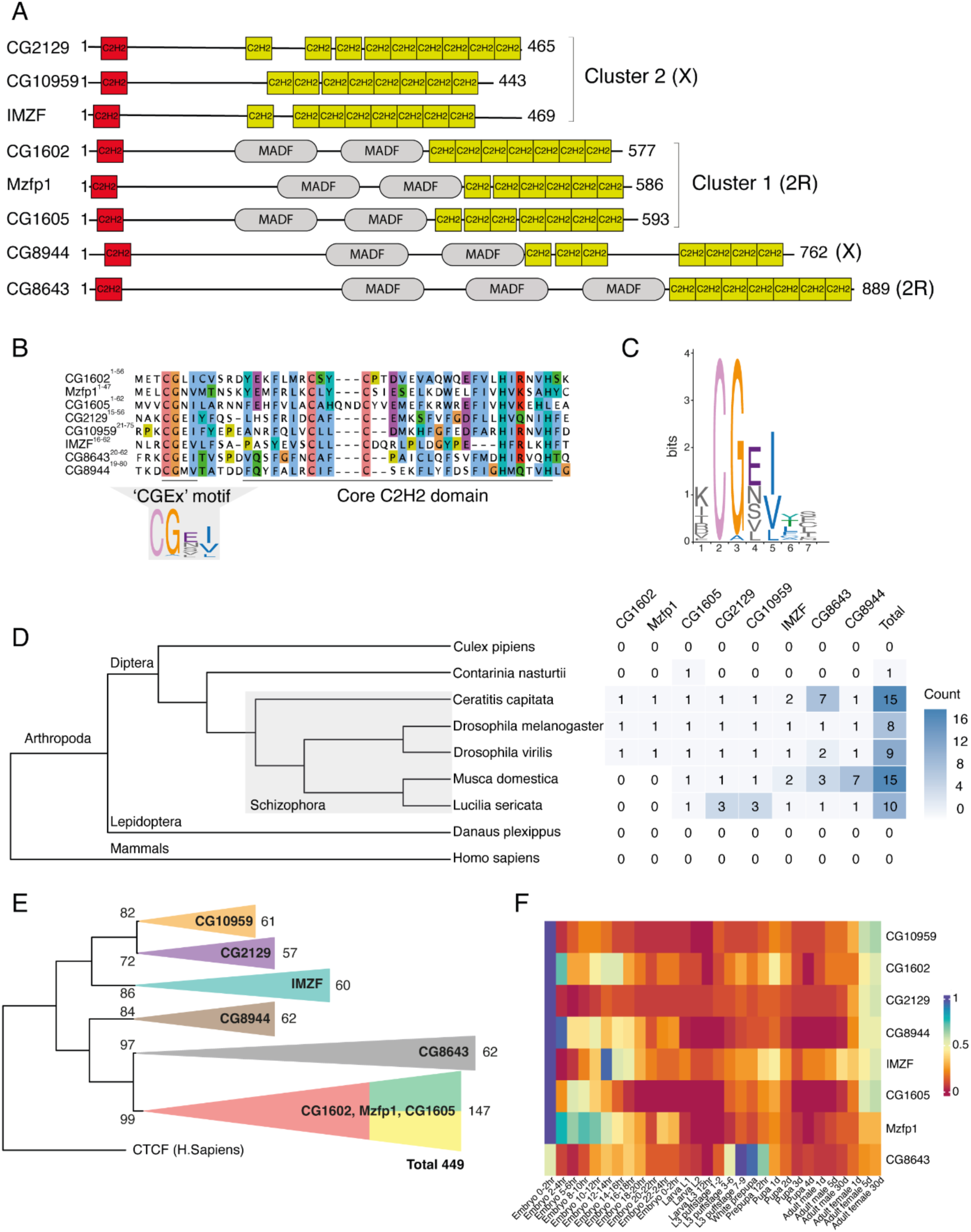
Domain architecture, phylogenetics and expression of’CGEx’-type C2H2 zinc finger-containing proteins. (**A**) Schematic representation of the domain structure of each studied C2H2 protein in *D. melanogaster*. UniProt (83) identifiers: CG2129 – Q9W3J4; CG10959 – Q9W3J2; IMZF – Q9W3J0; CG1602 – Q8MSB3; Mzfp1 – Q95TL6; CG1605 – Q5BIC3; CG8944 – Q9VXQ6; CG8643 – E1JH10. Chromosomal localization is indicated in round brackets. (**B**) Multiple sequence alignment of the N-terminal C2H2 domains of the studied proteins. Amino acid residues are colored with ClustalX (84) colors (blue – hydrophobic; red – positively charged; magenta – negatively charged; green – polar; pink – cysteines; orange – glycines; yellow – prolines; cyan – aromatic). Full ClustalX color scheme key is shown in **Supplementary Table S4**. (**C**) Hidden Markov Model of the’CGEx’ motif and its flanking amino acid residues. Residues are colored with ClustalX (84) colors (blue – hydrophobic; red – positively charged; magenta – negatively charged; green – polar; pink – cysteines; orange – glycines; yellow – prolines; cyan – aromatic). (**D**) Distribution of the studied C2H2 proteins among representative species, based on OrthoDB and EGGNOG data. For each species, the number of specific ortholog proteins is indicated on the heatmap. The phylogenetic relationships among taxa are based on the NCBI Taxonomy Database (66). (**E**) Schematic representation of the phylogenetic reconstruction of all analysed orthologs. The reconstruction was performed using IQ-TREE version 2 (74) with the maximum-likelihood (ML) method. Collapsed groups are color-coded as follows: CG10959 – orange; CG2129 – purple; IMZF – cyan; CG8944 – brown; CG8643 – black; CG1602, Mzfp1, and CG1605 were collapsed into a single group, represented by triangle divided into red, green, and yellow segments. Bootstrap values are indicated at the base of the nodes. The names of the orthologous groups are displayed inside the tree triangles, while the number of proteins in each group is shown outside. CTCF (CCCTC-binding factor) protein from *H. sapiens* (NCBI ID: 10664) was used as an outgroup during reconstruction (85). (**F**) Heatmap of the relative gene expression of C2H2 proteins throughout *D. melanogaster* development. Development stages are named according to FlyBase (78).

The genes encoding six out of eight proteins in the *Drosophila* genome reside within two gene clusters. Cluster 1, located on the 2R chromosome, includes genes for CG1602, Mzfp1, and CG1605 proteins, which, in addition to C2H2 zinc finger domains, encode a second type of DNA-binding domain known as MADF (myb/SANT-like domain in Adf-1) (81). Cluster 2, located on the X chromosome, comprises the non-MADF-containing genes for CG2129, CG10959, and the IMZF proteins. The genes for CG8643 (2R chromosome) and CG8944 (X chromosome) are located distantly from the described above clusters, both contain the MADF domains (**Figure 1A**).

Multiple sequence alignment (MSA) of these C2H2 domains revealed a high degree of similarity (**Figure 1B**). A distinguishing feature of these domains is a conserved N-terminal extension positioned upstream of the core C2H2 domain fold. Analysis of the MSA of orthologous proteins revealed that the characteristic N-terminal extension includes highly conserved cysteine, glycine, and glutamic acid residues, along with an additional hydrophobic residue (**Supplementary Figure S3**). Thus, we designated this region as the’CGEx’ motif, where x is a hydrophobic amino acid residue (**Figure 1B, 1C**). C2H2 domains containing this motif were classified as ‘CGEx’-type C2H2 domains.

Then we studied the evolutionary origin and prevalence of these domains across metazoan taxa. Analysis of orthologs of all eight proteins showed that’CGEx’-type C2H2 domains are restricted to a few Dipteran families, with the highest prevalence in Drosophilidae. Additional examples were identified in Glossinidae, Calliphoridae, Muscidae, Tephritidae, Diopsidae families (**Supplementary Table S3**), all of which are related to Schizophora (82). The most basal Diptera species in which’CGEx’-type C2H2 domain was identified is *Contarinia nasturtii* from Cecidomyiidae family. Notably, no proteins possessing’CGEx’-type C2H2 domains were detected in closely related families such as Culicidae or in other arthropods and mammals (**Figure 1D and Supplementary Table S3**). These findings indicate that’CGEx’-type C2H2 domains are a distinctive feature of Dipteran insects, enriched across diverse fly families.

Phylogenetic analysis of the orthologous proteins revealed strong, well-defined orthologous groups for each C2H2 protein (**Figure 1E and Supplementary Figure S4**). The colocalization of CG2129, CG10959, and IMZF within the same chromosomal cluster and phylogenetic branch, combined with their shared absence of MADF domains, suggests their common origin from a single ancestral gene via duplication. The subsequent step was the duplication of CG8944 on the X chromosome, in which MADF domains already appear. Further duplications gave rise to the CG8643, CG1602, Mzfp1, and CG1605 on chromosome 2R. Orthologs of CG1602, Mzfp1, and CG1605 formed a tight, unresolved branch due to high sequence similarity, indicative of recent gene duplications (**Figure 1E and Supplementary Figure S4**).

Analysis of the expression data revealed that family members exhibit their highest expression levels during the first two hours of *D. melanogaster* development, a period that corresponds to maternally deposited factors (**Figure 1F**).

### Dimers are the main oligomeric state of’CGEx’-type C2H2 domains

To test the homodimerization potential of all’CGEx’-type C2H2 domains from *D. melanogaster*, we performed yeast two-hybrid (Y2H) screening of the self-association ability of the polypeptides including single C2H2 domains with N-and C-terminal extensions. A complete list of the polypeptides is provided in **Supplementary Table S5**. Y2H assay showed homodimerization activity of 6 C2H2 domains – CG1602^1-56^, Mzfp1^1-47^, CG1605^1-62^, CG2129^1-56^, CG10959^1-75^, and IMZF^1-62^ (**Figure 2A** and **Supplementary Figure S5**). In the case of CG8944^1-80^ and CG8643^1-65^ the homodimerization activity was undetectable because of their strong self-activatory properties in Y2H assay (**Figure 2A**).

**Figure 2.**
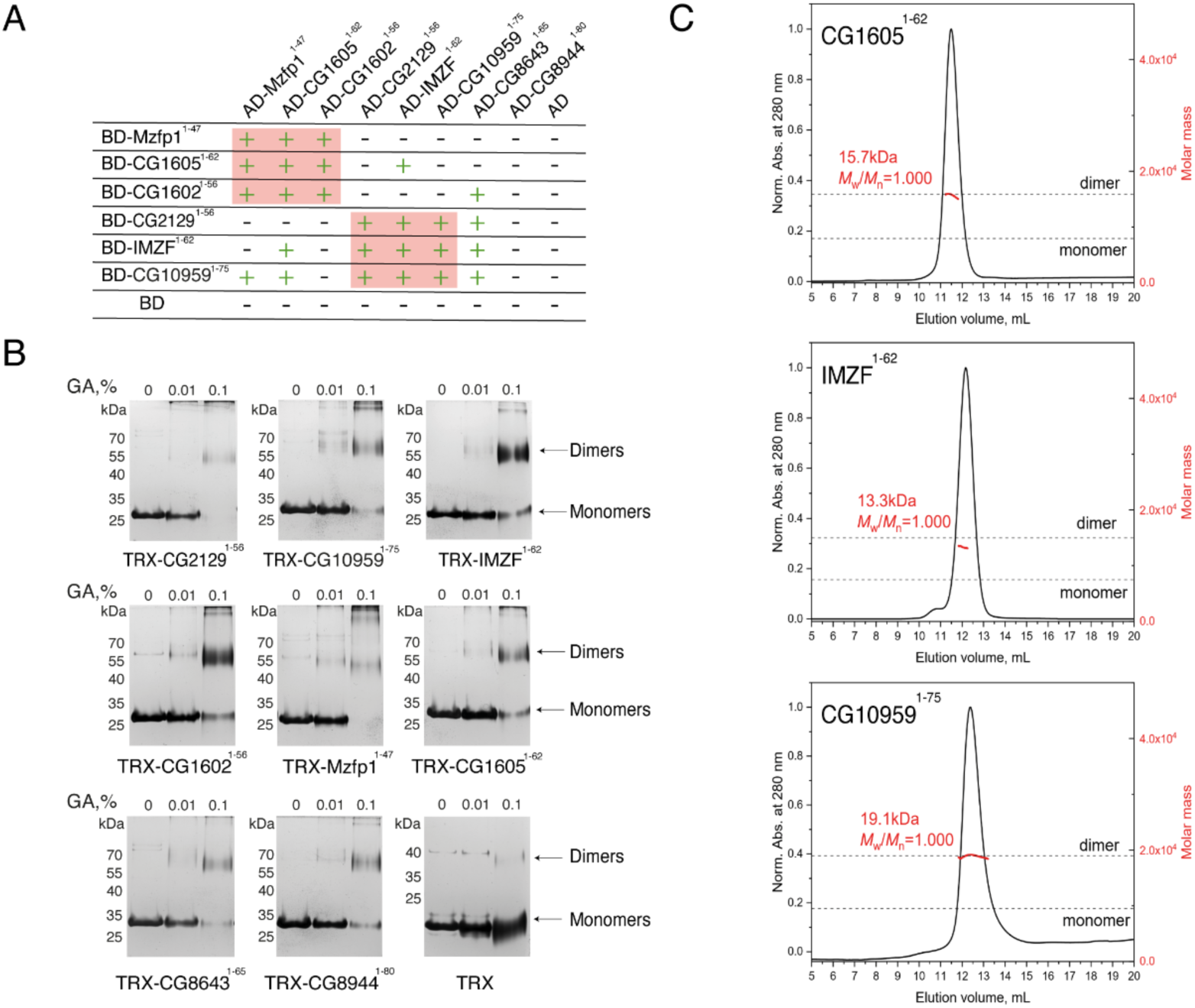
**Interactions and oligomeric state of’CGEx’-type C2H2 domains from *D. melanogaster* proteins.** (**A**) Yeast two-hybrid screening of the homo-and heterodimerization activities of ‘CGEx’-type C2H2 domains. Genome clusters of C2H2 proteins are highlighted with red squares. The activation domain (AD) was fused to the N-terminus of tested proteins. DNA-binding domain (BD) was fused to the N-or C-terminus of tested proteins (Mzfp^1-47^, CG2129^1-56^). Interaction is indicated by plus symbols; absence of interaction is marked with minus symbols. (**B**) Results of testing the homodimerization activity of each ‘CGEx’-type C2H2 domain using glutaraldehyde chemical cross-linking. Glutaraldehyde (GA) concentrations are indicated above the gel image. Molecular weight markers are shown on the left, and the observed oligomeric states are annotated on the right. (**C**) SEC-MALS analysis of wild-type CG1605^1-62^, IMZF^1-62^, and CG10959^1-75^ dimerization. MALS-derived molecular weights are indicated in red. Gray dashed lines mark the theoretical molecular weights of the monomers and dimers.

To confirm the homodimerization activity of the C2H2 domains *in vitro*, we performed glutaraldehyde chemical cross-linking for all studied C2H2 domains fused with thioredoxin as expression tag for better solubility. In all cases, we observed the disappearance of the band corresponding to monomers and the formation of higher molecular weight products corresponding to dimers (**Figure 2B**). For direct verification of the oligomerization state, we purified C2H2 domains of the IMZF^1-62^ (7.5 kDa), CG10959^1-75^ (9.5 kDa), and CG1605^1-62^ (7.9 kDa) proteins and determined their absolute molar mass using size-exclusion chromatography coupled with multi-angle light scattering (SEC-MALS). The elution profiles of all three purified C2H2 domains exhibited a single symmetrical peak, and SEC-MALS analysis confirmed that they exist as monodisperse homodimers: IMZF^1-62^ – 13.3 kDa, CG10959^1-75^ – 19.1 kDa, CG1605^1-62^ – 15.7 kDa (**Figure 2C**).

Given the sequence similarity of the’CGEx’-type C2H2 domains and their ability to form stable homodimers, we tested their propensity for heteromeric interactions using the Y2H assay (**Figure 2A** and **Supplementary Figure S5**). Our results showed that all family members are capable of forming heterodimers, primarily within their respective clusters. However, several interactions outside of these clusters were also observed. In particular, the C2H2 domain of IMZF^1-62^ interacted with CG1605^1-62^. The C2H2 domain of CG10959^1-75^, when fused toGAL4 DNA-binding domain (BD), formed heterodimers with Mzfp1^1-47^ and with CG1605^1-62^ fused to GAL4 activation domain (AD). The C2H2 domain of CG8643^1-65^, which does not belong to any cluster, demonstrated the ability to form heterodimers with proteins from both clusters. It interacted with CG1602^1-56^, CG2129^1-56^, and CG10959^1-75^, with a weak interaction also detected with IMZF^1-62^. Notably, no interaction was detected between CG8944^1-80^ and any other family members (**Figure 2A** and **Supplementary Figure S5**).

Overall,’CGEx’-type C2H2 domains exhibit a broad capacity for heteromeric interactions.

### The NMR structure of IMZF^1-62^ provides structural insights into the details of the molecular interface involved in single C2H2 domain dimer formation

Among the eight identified C2H2 domains, three — IMZF^1-62^, CG10959^1-75^, and CG1605^1-62^ — were successfully purified. Due to the failure of crystallization attempts, we selected IMZF^1-62^ for structural analysis by NMR spectroscopy, as it was obtained with the highest yield and purity.

The ^15^N HSQC spectrum of IMZF^1-62^ reveals a well-dispersed set of signals, indicative of a structured core, alongside regions of lower signal dispersion corresponding to flexible or unstructured fragments (**Figure 3A**, grey dots). The presence of a single set of resonances suggests that the protein exists as a symmetric homodimer in solution. Moreover, the absence of peak doubling or significant line broadening indicates a lack of detectable conformational heterogeneity, supporting the existence of a single, well-defined structural state.

**Figure 3.**
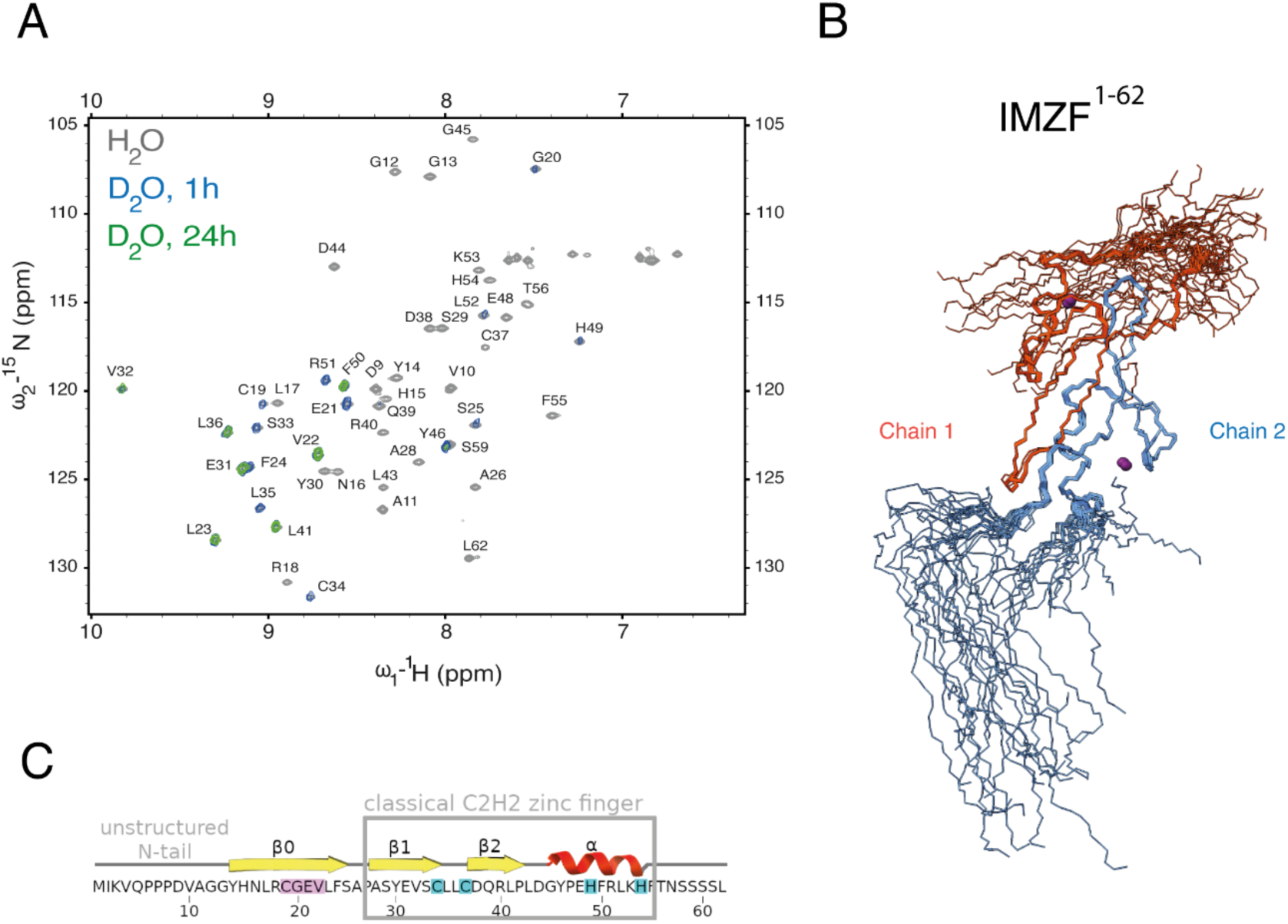
Structural characterization of ^15^N-labeled IMZF^1-62^ by NMR spectroscopy. (**A**) ^15^N–^1^H HSQC spectrum of ^15^N-IMZF^1-62^ acquired in 5% D₂O/95% H_2_O (grey dots) overlaid with spectra recorded after hydrogen/deuterium exchange in 100% D₂O for 1 hour (blue) and 24 hours (green). Backbone amide resonances are labeled. Spectra were recorded at 600 MHz (^1^H frequency) and 298 K in 20 mM sodium phosphate buffer (pH 7.0), containing 50 mM NaCl and 0.02% (w/v) sodium azide. (**B**) Superposition of the 20 lowest-energy NMR-derived structures of the IMZF^1-62^ dimer, shown with the chains colored distinctly. The ensemble members include the N-terminal extension (SGSPEF) remaining after TEV-protease cleavage. (**C**) Secondary structure of IMZF^1-62^, illustrating the classical zinc finger fold, an additional N-terminal β-strand (β0), and an unstructured N-terminal region. The conserved ‘CGEx’ motif is highlighted in pink; Zn-coordinating residues are highlighted in cyan.

The solution structure of IMZF^1-62^ was determined using a combination of NOE-derived distance restraints, torsion angle restraints, and hydrogen bond restraints (**Supplementary Tables S2A** and **S2B)**. Distance restraints between two chains were obtained from ^13^C/^15^N-filtered NOESY experiment, performed on a 1:2 mixture of isotopically labeled (^13^C/^15^N) and unlabeled protein. Residual dipolar couplings (RDCs) were not included in the structure determination, as the symmetric nature of the homodimer leads to opposing orientations of N–H bond vectors in each monomer, resulting in complete cancellation of RDCs.

The final ensemble of 20 NMR structures exhibits high convergence, with a backbone RMSD of 0.52 Å for the well-ordered regions (residues 13–55 of both monomers) upon superposition (**Figure 3B** and **Supplementary Table S2C**). The three-dimensional structure reveals that each monomer adopts a canonical C2H2 zinc finger fold, consisting of a β-hairpin followed by an α-helix, with a zinc ion coordinated by two cysteines and two histidines. Notably, the domain contains an additional N-terminal β-strand (residues 14–25), that includes the conserved’CGEx’ motif, as well as a disordered N-terminal tail (residues 1–13) extending beyond the structured core (**Figure 3C**).

The stability of the dimeric interface is further supported by hydrogen/deuterium exchange experiments, where numerous backbone amide protons remain protected from exchange even after 24 hours (**Figure 3A** and **Supplementary Figure S6**).

Quantitative analysis of backbone dynamics was performed through ^15^N relaxation measurements at 600 MHz (**Supplementary Figure S7**). The heteronuclear NOE values (panel A) and low order parameters (S² < 0.5, panel D) confirmed the disordered nature of the N-terminal tail (residues 1–13). In contrast, the core region encompassing the conserved’CGEx’ motif exhibited high order parameters (S² > 0.8, panel D), indicative of a rigid and well-structured conformation.

Elevated transverse relaxation rates (R₂, panel C) and substantial chemical exchange contributions (Rₑₓ > 2 s^-1^, panel E) were observed for residues at or near the dimer interface, suggesting micro-to millisecond timescale conformational exchange processes. These dynamic parameters were visually mapped onto the structural model in **Supplementary Figure S8**.

Rotational diffusion analysis of the dimer revealed pronounced anisotropy, with a diffusion tensor ratio (Dₐ/Dₓ) of approximately 2.0 (**Supplementary Figure S9**) and a mean correlation time (τ_c_) of 7.5 ns. The anisotropic diffusion model provided a markedly better fit to the experimental data than the isotropic approximation, consistent with the elongated, non-spherical geometry of the rigid dimeric core encompassing the’CGEx’ motif.

Dimer formation is stabilized by numerous inter-chain NOEs and hydrogen bonds. The interface is characterized by a striking six-stranded antiparallel β-sheet, formed through interactions between the additional β-strands (β0) of each monomer and their respective core β-strands (**Figure 4A**). The conserved residues C19, G20, E21, and V22 of the’CGEx’ motif constitute the structural core of the dimer interface, forming five symmetrical pairs of main-chain hydrogen bonds (**Figure 4A, 4B**). Specifically, two hydrogen bonds are formed between F24_chain1_ and L17_chain2,_ while V22_chain1_ engages in three hydrogen bonds — two with G20_chain2_ and one with C19_chain2_ residues. These interactions collectively generate an extensive hydrophobic interface of approximately 1080 Å² (**Figure 4C**).

**Figure 4.**
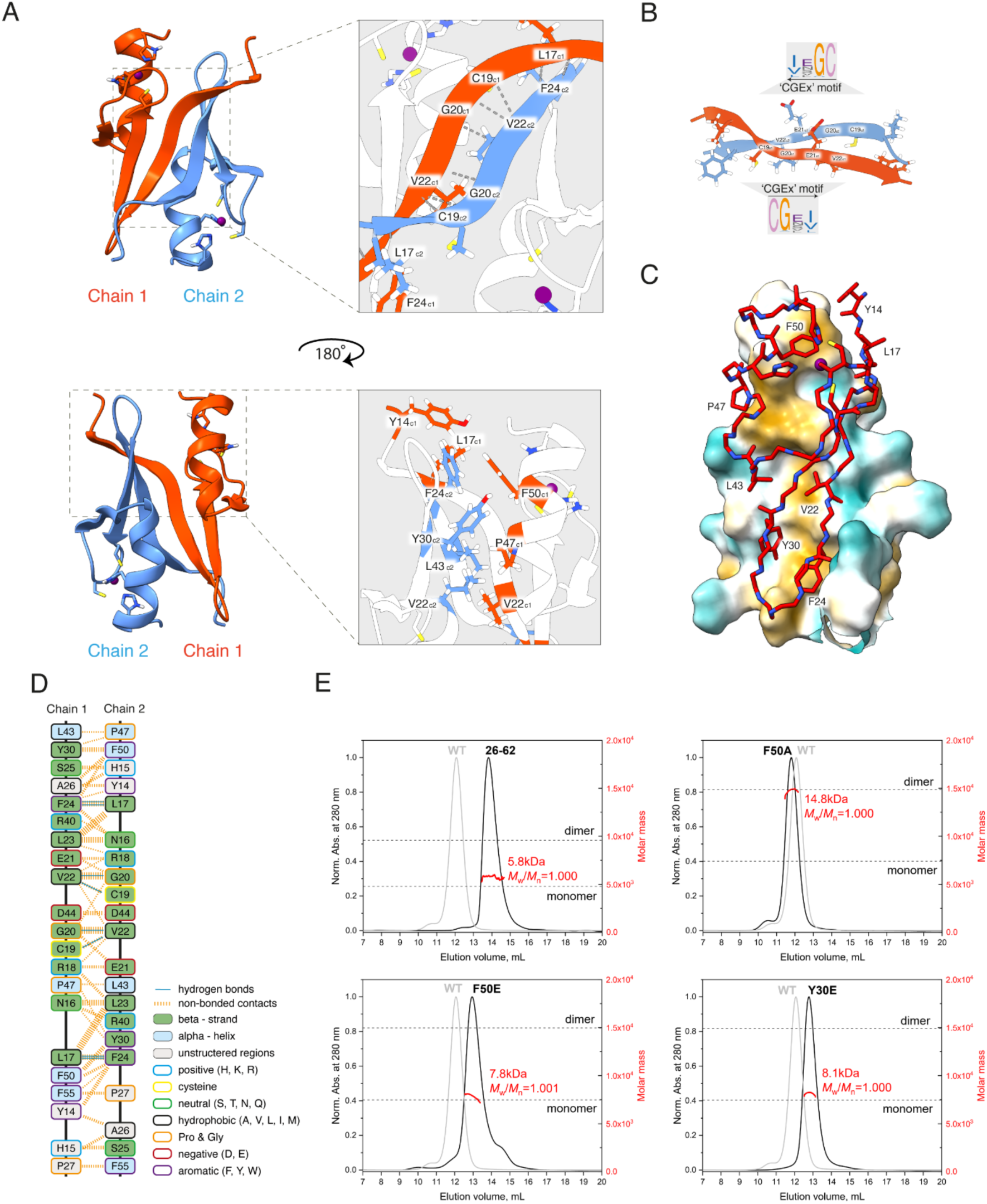
Structural and mutational analysis of the IMZF^1-62^ homodimer. (**A**) Representative structure of the IMZF^1-62^ homodimer highlighting key interacting residues. The structure is shown in cartoon representation, with each protein chain colored differently. Residues 14–55 are displayed. Hydrogen bonds are shown as grey lines. (**B**) Representative structure of the additional β-strands containing’CGEx’ motif. Hydrogen bonds are shown as grey lines. Amino acid residues are colored according to the ClustalX scheme based on HMM-profiles: blue – hydrophobic; red – positively charged; magenta – negatively charged; green – polar; pink – cysteines; orange – glycines; yellow – prolines; cyan – aromatic. The full ClustalX color key is provided in **Supplementary Table S4**. (**C**) Hydrophobic interface of the IMZF^1-62^ homodimer. Chain 1 is shown as a surface representation colored by hydrophobicity. Chain 2 is displayed as a red N-C**α**-C backbone trace. Nitrogen atoms are shown in blue; zinc ions are shown in purple. (**D**) Schematic representation of the IMZF^1-62^ dimer interface. Hydrogen bonds are indicated by blue lines; non-bonded contacts are shown as orange dashed lines. The width of the dashed line is proportional to the number of atomic contacts. The initial interacting scheme was generated using PDBsum (86). The color key is displayed to the right of the scheme. (**E)** SEC-MALS analysis of IMZF^1-62^ dimerization mutants. Elution profiles are labeled as WT (wild-type IMZF^1-62^), ‘26-62’ (truncated mutant lacking the’CGEx’ motif), and point mutants F50A, F50E, and Y30E. Shifts in elution volume reflect changes in oligomeric state. MALS-derived molecular weights are indicated in red. Gray dashed lines mark the theoretical molecular weights of the monomers and dimers.

Symmetric hydrophobic contacts are concentrated within a compact hydrophobic groove on the surface of the C2H2 core, involving residues from both the α-helix and β-hairpin (**Figure 4A, 4C**). Key residues from one monomer that insert into this groove include Y14, L17, V22, F24, Y30, L43, P47, and F50. The side-chain of F24_chain1_ engages in extensive hydrophobic interactions with side chains of Y14_chain2_, F50_chain2_, and L17_chain2_, while Y30_chain1_ also contacts F50_chain2_. Additionally, L43_chain1_ interacts with the P47_chain2_ via side-chains contact. A detailed map of the dimer interface is provided in **Figure 4D**.

Due to the high sequence similarity among all identified C2H2 domains, they are predicted to adopt the same overall structure as IMZF^1-62^ (IMZF). AlphaFold2 confidently predicted a conserved dimeric architecture for all eight ‘CGEx’ motif-containing C2H2 domains, with models that are readily superimposable onto the experimentally determined NMR structure of IMZF^1-62^ (**Supplementary Figure S10**).

According to these predictions, each domain features an additional β-strand located upstream of the canonical zinc finger core, and conserves key amino acid residues involved in hydrophobic packing within the dimer interface. Overall, the predicted structures exhibit high structural similarity to the IMZF^1-62^ dimer.

To investigate the mechanism of dimer formation and clarify the role of the’CGEx’ motif, we performed site-directed mutagenesis of IMZF^1-62^. Given the contribution of the’CGEx’ motif to homodimerization, we purified a truncated variant, IMZF^26-62^ (∼4.9 kDa), lacking this motif, and determined its absolute molar mass using SEC-MALS. Size-exclusion chromatography revealed a single, symmetric peak that eluted later than the IMZF^1-62^ dimer, while MALS analysis yielded a molecular mass of 5.8 kDa (∼1.3-mer), consistent with substantial monomerization. These results underscore the essential role of the’CGEx’ motif in stabilizing the dimeric assembly (**Figure 4E**).

Next, guided by the NMR structure of IMZF^1-62^, we introduced single-point mutations at residues involved in key hydrophobic interactions within the core of the C2H2 domain. Specifically, Y30 and F50 were replaced with glutamate (Y30E and F50E) to disrupt hydrophobic packing. In both mutants, SEC-MALS analysis revealed single symmetrical peaks shifted relative to that of the IMZF^1-62^ dimer, with molecular masses consistent with monomeric species: 8.1 kDa for Y30E and 7.8 kDa for F50E (**Figure 4E** and **Supplementary Table S6**).

In contrast, substitution of F50 with alanine (F50A) did not disrupt dimerization. The MALS-derived molecular weight for IMZF^1-62^ F50A was 14.8 kDa, consistent with a dimeric state (**Figure 4E** and **Supplementary Table S6**). Together, these mutational data demonstrate that dimerization depends on both the conserved’CGEx’ motif and specific hydrophobic interactions within the C2H2 core.

### The presence of additional secondary structure elements adjacent to C2H2 domains is a widespread phenomenon and is most prevalent in insects

To study the distribution of C2H2 domains with additional secondary structure elements beyond the core fold in the proteomes of various organisms, we developed an algorithm implemented in Python. The first step is the analysis of the secondary structure of C2H2 proteins from whole-proteome databases of protein structure models obtained using AlphaFold2 (79). Then, domains annotated as C2H2 zinc fingers are examined for the presence of additional secondary structure elements within a 20-residue window preceding the core C2H2 fold, provided there are no other annotated domains in this region. Then, amino acid sequences of domains with β-strand extensions were exported and the structures of their dimers were predicted using AlphaFold2-Multimer (87). Structures with interchain PAE score below 5.0 and pLDDT values of residues at the predicted dimerization interface above 70 were further inspected visually.

The analysis of 27 proteomes of living organisms, representing distinct phylogenetic groups of higher metazoans, revealed that dimerization mediated by C2H2 domains with additional structural elements is a common phenomenon (**Figure 5A** and **Supplementary Table S7**). Structural elements similar to those found in IMZF orthologs (‘CGEx’ motif) were observed exclusively in insects, for which several other types of dimeric assemblies are also predominantly found. We also detected one C2H2 domain in *Octopus bimaculoides* (UniProt ID: A0A0L8FQ77) and two domains in *Acyrthosiphon pisum* (UniProt IDs: J9LEP9, X1X1C4) that adopt an IMZF-like fold. However, these domains do not contain the ‘CGEx’ motif and are not classified as orthologs according to orthology databases, indicating that the observed fold likely arose through convergent evolution (**Figure 5A** and **Supplementary Table S7**).

**Figure 5.**
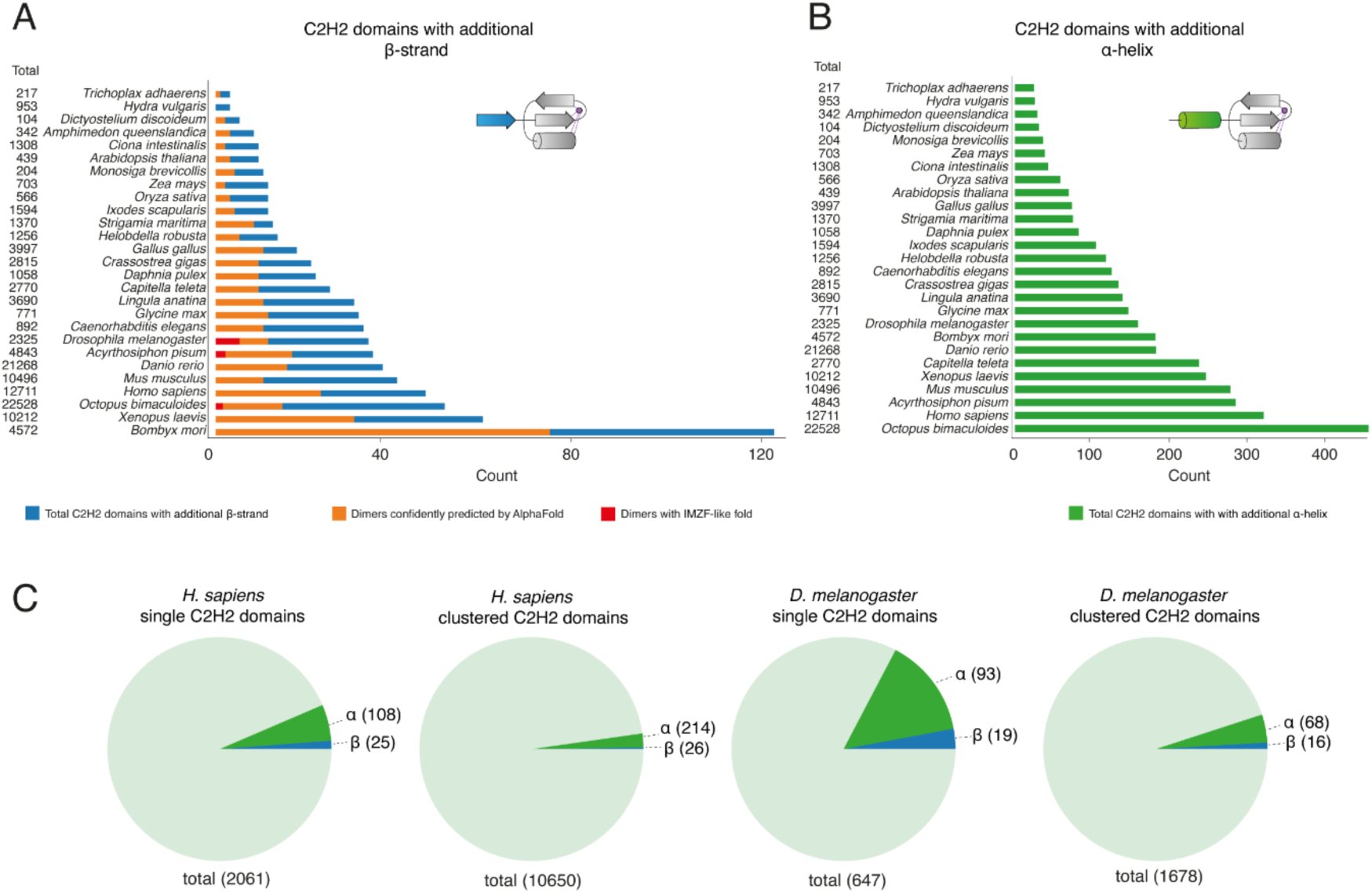
The presence of C2H2 domains with additional N-terminal secondary structure elements across the proteomes of higher eukaryotes. (**A**) The distribution of C2H2 zinc fingers with an additional N-terminal β-strand across various species of multicellular eukaryotes. The total numbers of predicted C2H2 domains in the proteome of each species are shown to the left of the corresponding species name. Blue bars represent the total number of predicted C2H2 zinc fingers containing an additional β-strand. Orange bars show the subset of those C2H2 domains predicted by AlphaFold2 to form confident dimers. Red squares mark C2H2 domains adopting an IMZF-like dimeric fold. (**B**) The distribution of C2H2 zinc fingers with an additional N-terminal α-helix by species. Green bars show the number of C2H2 domains predicted to contain an additional α-helix. The total numbers of C2H2 domains in each proteome are shown on the left. (**C**) The relative abundance of C2H2 domains with additional secondary structure elements (within the window of 20-residues preceding the core C2H2 fold with the absence of other annotated domains spanning within this window) among single and clustered (10 or fewer residues between domains) C2H2 domains in the genomes of *H. sapiens* and *D. melanogaster*. Total numbers of C2H2 zinc fingers are shown below. α – the number of C2H2 domains that possess an additional α-helix. β – the number of C2H2 zinc fingers possessing an additional β-strand. Further details are shown in **Supplementary Table S7**.

Another group of C2H2-type domains feature an additional α-helix at the N-terminus. According to AlphaFold2 predictions for some representatives, these α-helices can participate in heteromeric protein–protein interactions. However, prediction of their tentative dimerization activity did not produce reliable results. The total number of C2H2 domains containing an additional α-helix is quite large in all studied proteomes (**Figure 5B**). Noteworthy, the AlphaFold database contains predictions only for monomeric assemblies, but some sequences typically are not predicted to form secondary structure in the absence of an interaction partner, thus the overall quantity of domains with additional elements is likely to be underestimated.

We then inspected the relative abundance of C2H2 domains with adjacent secondary structure elements among single and clustered (10 or less residues between individual domains) C2H2 domain populations (**Figure 5C**). In general, C2H2 domains with adjacent secondary structure elements are more frequent among single zinc fingers. This further supports their higher propensity for protein–protein interaction rather than DNA binding, which is more characteristic of clustered C2H2 domains.

Overall, the presence of additional secondary structure elements is a widespread feature of C2H2 domains. In many cases, especially in insects, an additional β-strand provides the potential for the domain to gain dimerization capacity.

## DISCUSSION

In this study, we characterized an eight-member C2H2 protein family in *Drosophila melanogaster*. Each of these proteins contains an N-terminal C2H2 domain capable of forming stable dimers. The NMR structure of the dimeric C2H2 domain from the IMZF protein revealed an additional β-strand bearing a conserved’CGEx’ motif, which is essential for dimerization. The dimer interface also involves the core β-strands and the α-helix of the C2H2 domain. Notably, residues from the α-helix interact with both the core β-strands and the β-strand containing the’CGEx’ motif. In contrast, the canonical DNA-binding residues at positions-1, 2, 3, and 6 relative to the α-helix do not contribute to dimer formation (**Figures 3C, 4A, 4B**) (11, 88). Although few structural examples of direct interaction between zinc fingers are available, existing data indicate that these domains can utilize diverse amino acid positions to mediate inter-domain contacts. For instance, the C2H2 protein ZAP1 contains tandem zinc fingers that interact via their α-helices, without involving canonical DNA-binding residues (34). In contrast, the atypical C4 zinc finger of GATA-1 and the C3H zinc finger of the FOG protein form a complex via α-helical contacts that include DNA-binding positions (**Figure 6**) (44).

**Figure 6.**
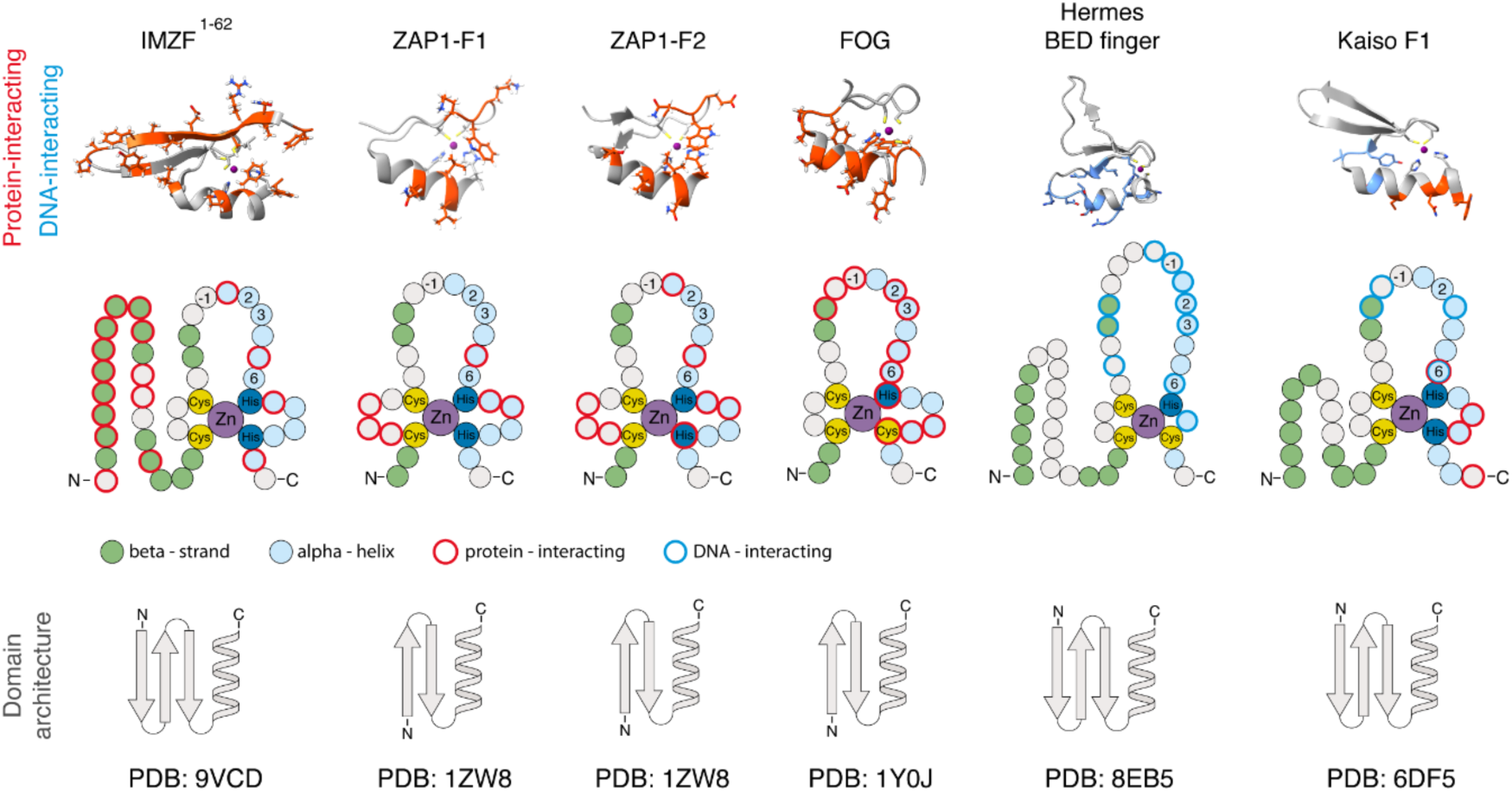
Comparison of zinc finger residues used for interaction with other zinc fingers and DNA. Cartoons were drawn according to the following structures (PDB ID): 9VCD (IMZF^1-62^), 1ZW8 (ZAP1-Finger 1 and ZAP1-Finger 2) (34), 1Y0J (FOG-Finger 1 and GATA) (44), 8EB5 (Hermes BED finger) (90), 6DF5 (Kaiso-Finger 1) (94).

A distinguishing feature of the IMZF^1-62^ domain is its use of an additional secondary structure element in dimerization. Few zinc finger domains containing extra β-strands have been described to date. For example, BED fingers, found in various proteins in *D. melanogaster* and *H. sapiens*, include an additional N-terminal β-strand, although its functional significance remains unclear. In contrast to the dimerizing C2H2 domain of IMZF, BED fingers are known to function exclusively in DNA binding and employ canonical amino acid residues within the α-helix to recognize DNA (**Figure 6**) (89, 90).

The Ikaros family of transcription factors provides a well-characterized example of direct interactions mediated by C2H2 zinc fingers. The Ikaros multimerization domain comprises double C2H2 motifs located within the same polypeptide chain (40, 91), both of which are required for dimerization. This mechanism apparently differs fundamentally from that observed in single ‘CGEx’-type C2H2 domains and remains to be fully elucidated.

Our results demonstrate the broad capacity of the studied C2H2 domains to engage in heterotypic interactions, as revealed by Y2H assay. These domains predominantly form heterodimers within their respective gene clusters, although several inter-cluster interactions were also detected. Unexpectedly, we did not detect heterodimer formation involving the C2H2 domain of CG8944 with any other C2H2 domain tested. This suggests that CG8944 may have undergone rapid evolutionary divergence and possesses unique structural features that prevent heterodimerization. Importantly, heterodimer formation via the C2H2 effector domain is likely subject to strict regulation—possibly co-translationally or through other cellular mechanisms— with homodimer formation being the preferred mode of interaction (92, 93). ‘CGEx’-type C2H2 domains appear to be a unique feature of the Schizophora group of the Diptera order, as no orthologous proteins containing this type of zinc finger were identified in other taxa. This allows us to assume that the primary expansion of the corresponding genes occurred already in the common ancestor of Schizophora or slightly earlier, probably during the last period of mass diversification of Diptera (82). The most distantly related species to Schizophora in the Diptera order, which has a single orthologous protein (NCBI ID: 116339888) containing the ‘CGEx’-type C2H2 domain, is *Contarinia nasturtii* from the Cecidomyiidae family. This ortholog, which lacks MADF domains and occupies a basal position in the phylogenetic tree, may represent an ancestral form for all other ‘CGEx’-type C2H2 domain-containing proteins. Therefore, the primary duplication that led to the formation of the most ancient cluster 2 (CG2129, CG10959, IMZF) likely occurred between the divergence of the infraorder Bibionomorpha and the radiation of Schizophora, ∼250–75 million years ago, according to Wiegmann et al. (82).

Analysis of the presence of additional secondary structure elements revealed that this feature is widespread across proteomes of higher metazoans. Protein modelling suggests that some of these domains can possess dimerization activity, whereas the mechanisms of dimer formation may differ. The C2H2 domain dimerization has likely arisen independently several times during evolution, since domains from phylogenetically distant groups typically do not show significant amino acid sequence similarity and often have dissimilar predicted structures. Obviously, functions of additional N-terminal β-strands are not limited only to dimerization; they may participate in other protein–protein interactions and in DNA binding, as observed for the first zinc finger of human Kaiso protein (8). C2H2 domains bearing α-helices or β-strands preceding the core fold are more abundant among single than clustered zinc fingers further highlighting their likely engagement in protein–protein interaction. Earlier studies have suggested that IMZF and Mzfp1 function as transcriptional regulators. IMZF is involved in the immune response (47), while Mzfp1 exhibits architectural activity (45) and plays a role in regulating the MAPK pathway (46). It is likely that other members of this family also function as architectural proteins and regulators of gene expression. Expression profiling revealed that family members are most highly expressed during early *D. melanogaster* development, suggesting a regulatory role for these C2H2 proteins in early developmental processes. Dimerization of’CGEx’-type zinc finger domains in this group of C2H2 proteins may play a role in increasing the specificity and efficiency of recruitment to regulatory elements and organizing long-distance interactions between their binding sites of these proteins. The functional role of dimerization is a subject for future research.

## ACKNOWLEDGMENTS

The authors are grateful to V. Sokolov for assistance with molecular cloning.

This study was performed using the equipment of the IGB RAS Core Facilities Centre.

## AUTHOR CONTRIBUTIONS

K.I.B. designed the research studies, supervised the experiments, conducted molecular cloning, protein expression and purification, yeast two-hybrid assays, and bioinformatic analysis; prepared figures; and drafted, wrote and revised the manuscript. S.S.M. supervised, conducted and analysed NMR experiments, wrote, and revised the manuscript. S.D.D. analysed NMR experiments. N.N.S. conducted SEC-MALS experiments and revised the manuscript. A.M.K. supervised the bioinformatic analysis and revised the manuscript. O.G.M. supervised the research and revised the manuscript. A.A.G. conducted protein expression and purification and revised the manuscript. O.V.A. conducted protein expression and purification. A.A.S. conducted protein expression and purification and chemical cross-linking assays.

E.V.B. conducted NMR experiments. V.I.P. supervised NMR experiments and revised the manuscript. P.G.G. supervised the experiments and revised the manuscript.

A.N.B. initiated the project, designed research studies, conducted bioinformatic analysis, supervised the experiments, and wrote and revised the manuscript.

## SUPPLEMENTARY DATA

Supplementary Data are available at NAR online.

## DATA AVAILABILITY

The final NMR ensemble has been deposited in the Protein Data Bank (PDB ID: 9VCD), and the corresponding chemical shift assignments are available in the BioMagResBank (BMRB-36762).

The in-house built Python scripts used to analysis of the proteome-wide presence of additional structure elements of C2H2 domains using DSSP algorithm (95) (based on AlphaFold2 predictions) are available at GitHub (https://github.com/errinaceus/DSSP_evaluation).

The in-house built R scripts used for phylogenetic analysis are available at GitHub (https://github.com/kbalagurov/IMZF-CG18262_orthologs_analysis).

## REFERENCES

1. Iuchi, S. (2001) Three classes of C2H2 zinc finger proteins: *CMLS*, Cell. Mol. Life Sci., 58, 625–635. 10.1007/PL00000885

2. Bonchuk, A.N. and Georgiev, P.G. (2024) C2H2 proteins: Evolutionary aspects of domain architecture and diversification. BioEssays, 46, 2400052. 10.1002/bies.202400052

3. Fedotova, A.A., Bonchuk, A.N., Mogila, V.A., et al. (2017) C2H2 zinc finger proteins: the largest but poorly explored family of higher eukaryotic transcription factors. Acta Naturae, 9, 47–58. 10.32607/20758251-2017-9-2-47-58

4. Miller, J., McLachlan, A.D. and Klug, A. (1985) Repetitive zinc-binding domains in the protein transcription factor IIIA from Xenopus oocytes. EMBO J, 4, 1609–1614. 10.1002/j.1460-2075.1985.tb03825.x

5. Krishna, S.S., Majumdar, I. and Grishin, N.V. (2003) Structural classification of zinc fingers: survey and summary. Nucleic Acids Res, 31, 532–550. 10.1093/nar/gkg161

6. Matthews, J.M. and Sunde, M. (2002) Zinc fingers--folds for many occasions. IUBMB Life, 54, 351–355. 10.1080/15216540216035

7. Babu, M.M., Iyer, L.M., Balaji, S., et al. (2006) The natural history of the WRKY– GCM1 zinc fingers and the relationship between transcription factors and transposons. Nucleic Acids Research, 34, 6505–6520. 10.1093/nar/gkl888

8. Buck-Koehntop, B.A., Stanfield, R.L., Ekiert, D.C., et al. (2012) Molecular basis for recognition of methylated and specific DNA sequences by the zinc finger protein Kaiso. Proc. Natl. Acad. Sci. U.S.A., 109, 15229–15234. 10.1073/pnas.1213726109

9. Foster, M.P., Wuttke, D.S., Radhakrishnan, I., et al. (1997) Domain packing and dynamics in the DNA complex of the N-terminal zinc fingers of TFIIIA. Nat Struct Biol, 4, 605–608. 10.1038/nsb0897-605

10. Pavletich, N.P. and Pabo, C.O. (1991) Zinc finger-DNA recognition: crystal structure of a Zif268-DNA complex at 2.1 Å. Science, 252, 809–817. 10.1126/science.2028256

11. Wolfe, S.A., Nekludova, L. and Pabo, C.O. (2000) DNA Recognition by Cys_2_His_2_ Zinc Finger Proteins. Annu. Rev. Biophys. Biomol. Struct., 29, 183–212. 10.1146/annurev.biophys.29.1.183

12. Maksimenko, O.G., Fursenko, D.V., Belova, E.V., et al. (2021) CTCF as an example of DNA-binding transcription factors containing clusters of C2H2-type zinc fingers. Acta Naturae, 13, 31–46. 10.32607/actanaturae.11206

13. Kyrchanova, O.V., Bylino, O.V. and Georgiev, P.G. (2022) Mechanisms of enhancer-promoter communication and chromosomal architecture in mammals and *Drosophila*. Front. Genet., 13, 1081088. 10.3389/fgene.2022.1081088

14. Kyrchanova, O., Sokolov, V. and Georgiev, P. (2023) Mechanisms of interaction between enhancers and promoters in three *Drosophila* model systems. IJMS, 24, 2855. 10.3390/ijms24032855

15. Bonchuk, A., Balagurov, K. and Georgiev, P. (2023) BTB domains: A structural view of evolution, multimerization, and protein–protein interactions. BioEssays, 45, 2200179. 10.1002/bies.202200179

16. Olivieri, D., Paramanathan, S., Bardet, A.F., et al. (2021) The BTB-domain transcription factor ZBTB2 recruits chromatin remodelers and a histone chaperone during the exit from pluripotency. Journal of Biological Chemistry, 297, 100947. 10.1016/j.jbc.2021.100947

17. Bonchuk, A.N., Balagurov, K.I., Baradaran, R., et al. (2024) The Arthropoda-specific Tramtrack group BTB protein domains use previously unknown interface to form hexamers. eLife, 13, e96832. 10.7554/eLife.96832

18. Bonchuk, A., Denisov, S., Georgiev, P., et al. (2011) *Drosophila* BTB/POZ domains of ‘ttk group’ can form multimers and selectively interact with each other. Journal of Molecular Biology, 412, 423–436. 10.1016/j.jmb.2011.07.052

19. Huynh, K.D. and Bardwell, V. J. (1998) The BCL-6 POZ domain and other POZ domains interact with the co-repressors N-CoR and SMRT. Oncogene, 17, 2473– 2484. 10.1038/sj.onc.1202197

20. Mance, L., Bigot, N., Zhamungui Sánchez, E., et al. (2024) Dynamic BTB-domain filaments promote clustering of ZBTB proteins. Molecular Cell, 84, 2490–2510.e9. 10.1016/j.molcel.2024.05.029

21. Maksimenko, O., Kyrchanova, O., Klimenko, N., et al. (2020) Small *Drosophila* zinc finger C2H2 protein with an N-terminal zinc finger-associated domain demonstrates the architecture functions. Biochim Biophys Acta Gene Regul Mech, 1863, 194446. 10.1016/j.bbagrm.2019.194446

22. Bonchuk, A., Boyko, K., Fedotova, A., et al. (2021) Structural basis of diversity and homodimerization specificity of zinc-finger-associated domains in *Drosophila*. Nucleic Acids Research, 49, 2375–2389. 10.1093/nar/gkab061

23. Schumacher, C., Wang, H., Honer, C., et al. (2000) The SCAN domain mediates selective oligomerization. Journal of Biological Chemistry, 275, 17173–17179. 10.1074/jbc.M000119200

24. Ecco, G., Imbeault, M. and Trono, D. (2017) KRAB zinc finger proteins. Development, 144, 2719–2729. 10.1242/dev.132605

25. Collins, T., Stone, J.R. and Williams, A.J. (2001) All in the family: the BTB/POZ, KRAB, and SCAN domains. Molecular and Cellular Biology, 21, 3609–3615. 10.1128/MCB.21.11.3609-3615.2001

26. Frietze, S. and Farnham, P.J. (2011) Transcription factor effector domains. Subcell Biochem, 52, 261–277. 10.1007/978-90-481-9069-0_12

27. Datta, R.R. and Rister, J. (2022) The power of the (imperfect) palindrome: Sequence-specific roles of palindromic motifs in gene regulation. BioEssays, 44, 2100191. 10.1002/bies.202100191

28. Funnell, A.P.W. and Crossley, M. (2012) Homo-and heterodimerization in transcriptional regulation. In Matthews, J.M. (ed), Protein Dimerization and Oligomerization in Biology, Advances in Experimental Medicine and Biology. Springer New York, New York, NY, Vol. 747, pp. 105–121. 10.1007/978-1-4614-3229-6_7

29. Ortega, E., Rengachari, S., Ibrahim, Z., et al. (2018) Transcription factor dimerization activates the p300 acetyltransferase. Nature, 562, 538–544. 10.1038/s41586-018-0621-1

30. Gamsjaeger, R., Liew, C., Loughlin, F., et al. (2007) Sticky fingers: zinc-fingers as protein-recognition motifs. Trends in Biochemical Sciences, 32, 63–70. 10.1016/j.tibs.2006.12.007

31. Brayer, K.J. and Segal, D.J. (2008) Keep your fingers off my DNA: protein–protein interactions mediated by C2H2 zinc finger domains. Cell Biochem Biophys, 50, 111– 131. 10.1007/s12013-008-9008-5

32. Simpson, R.J.Y., Yi Lee, S.H., Bartle, N., et al. (2004) A classic zinc finger from friend of GATA mediates an interaction with the coiled-coil of transforming acidic coiled-coil 3. Journal of Biological Chemistry, 279, 39789–39797. 10.1074/jbc.M404130200

33. Omichinski, J.G., Clore, G.M., Robien, M., et al. (1992) High-resolution solution structure of the double Cys2His2 zinc finger from the human enhancer binding protein MBP-1. Biochemistry, 31, 3907–3917. 10.1021/bi00131a004

34. Wang, Z., Feng, L.S., Matskevich, V., et al. (2006) Solution structure of a Zap1 zinc-responsive domain provides insights into metalloregulatory transcriptional repression in *Saccharomyces cerevisiae*. Journal of Molecular Biology, 357, 1167– 1183. 10.1016/j.jmb.2006.01.010

35. Wojtaszek, J.L., Wang, S., Kim, H., et al. (2014) Ubiquitin recognition by FAAP20 expands the complex interface beyond the canonical UBZ domain. Nucleic Acids Research, 42, 13997–14005. 10.1093/nar/gku1153

36. Suzuki, N., Rohaim, A., Kato, R., et al. (2016) A novel mode of ubiquitin recognition by the ubiquitin-binding zinc finger domain of WRNIP 1. The FEBS Journal, 283, 2004–2017. 10.1111/febs.13734

37. Kastner, P., Aukenova, A. and Chan, S. (2024) Evolution of the Ikaros family transcription factors: From a deuterostome ancestor to humans. Biochemical and Biophysical Research Communications, 694, 149399. 10.1016/j.bbrc.2023.149399

38. Kelley, C.M., Ikeda, T., Koipally, J., et al. (1998) Helios, a novel dimerization partner of Ikaros expressed in the earliest hematopoietic progenitors. Current Biology, 8, 508–S1. 10.1016/S0960-9822(98)70202-7

39. Perdomo, J., Holmes, M., Chong, B., et al. (2000) Eos and Pegasus, two members of the Ikaros family of proteins with distinct DNA binding activities. Journal of Biological Chemistry, 275, 38347–38354. 10.1074/jbc.M005457200

40. McCarty, A.S., Kleiger, G., Eisenberg, D., et al. (2003) Selective dimerization of a C2H2 zinc finger subfamily. Molecular Cell, 11, 459–470. 10.1016/S1097-2765(03)00043-1

41. Morgan, B., Sun, L., Avitahl, N., et al. (1997) Aiolos, a lymphoid restricted transcription factor that interacts with Ikaros to regulate lymphocyte differentiation. EMBO J, 16, 2004–2013. 10.1093/emboj/16.8.2004

42. Westman, B.J., Perdomo, J., Sunde, M., et al. (2003) The C-terminal domain of Eos forms a high order complex in solution. Journal of Biological Chemistry, 278, 42419–42426. 10.1074/jbc.M306817200

43. Sun, L., Liu, A. and Georgopoulos, K. (1996) Zinc finger-mediated protein interactions modulate Ikaros activity, a molecular control of lymphocyte development. The EMBO Journal, 15, 5358–5369. 10.1002/j.1460-2075.1996.tb00920.x

44. Liew, C.K., Simpson, R.J.Y., Kwan, A.H.Y., et al. (2005) Zinc fingers as protein recognition motifs: structural basis for the GATA-1/friend of GATA interaction. Proc Natl Acad Sci U S A, 102, 583–588. 10.1073/pnas.0407511102

45. Sokolov, V., Kyrchanova, O., Klimenko, N., et al. (2024) New *Drosophila* promoter-associated architectural protein Mzfp1 interacts with CP190 and is required for housekeeping gene expression and insulator activity. Nucleic Acids Research, 52, 6886–6905. 10.1093/nar/gkae393

46. Ashton-Beaucage, D., Udell, C.M., Gendron, P., et al. (2014) A functional screen reveals an extensive layer of transcriptional and splicing control underlying RAS/MAPK signaling in Drosophila. PLoS Biol, 12, e1001809. 10.1371/journal.pbio.1001809

47. Zhou, H., Liu, L., Pang, Y., et al. (2024) Relish-mediated C2H2 zinc finger protein IMZF restores *Drosophila* immune homeostasis via inhibiting the transcription of *Imd/Tak1*. Insect Biochemistry and Molecular Biology, 170, 104138. 10.1016/j.ibmb.2024.104138

48. Marley, J., Lu, M. and Bracken, C. (2001) A method for efficient isotopic labeling of recombinant proteins. J Biomol NMR, 20, 71–75. 10.1023/a:1011254402785

49. Delaglio, F., Grzesiek, S., Vuister, G.W., et al. (1995) NMRPipe: a multidimensional spectral processing system based on UNIX pipes. J Biomol NMR, 6, 277–293. 10.1007/BF00197809

50. Lee, W., Rahimi, M., Lee, Y., et al. (2021) POKY: a software suite for multidimensional NMR and 3D structure calculation of biomolecules. Bioinformatics, 37, 3041–3042. 10.1093/bioinformatics/btab180

51. Bahrami, A., Assadi, A.H., Markley, J.L., et al. (2009) Probabilistic interaction network of evidence algorithm and its application to complete labeling of peak lists from protein NMR spectroscopy. PLoS Comput Biol, 5, e1000307. 10.1371/journal.pcbi.1000307

52. Polshakov, V.I., Frenkiel, T.A., Birdsall, B., et al. (1995) Determination of stereospecific assignments, torsion-angle constraints, and rotamer populations in proteins using the program AngleSearch. J Magn Reson B, 108, 31–43. 10.1006/jmrb.1995.1099

53. Shen, Y. and Bax, A. (2013) Protein backbone and sidechain torsion angles predicted from NMR chemical shifts using artificial neural networks. J Biomol NMR, 56, 227–241. 10.1007/s10858-013-9741-y

54. Bardiaux, B., Malliavin, T. and Nilges, M. (2012) ARIA for solution and solid-state NMR. In Shekhtman, A., Burz, D.S. (eds), Protein NMR Techniques, Methods in Molecular Biology. Humana Press, Totowa, NJ, Vol. 831, pp. 453–483. 10.1007/978-1-61779-480-3_23

55. Istrate, A.N., Tsvetkov, P.O., Mantsyzov, A.B., et al. (2012) NMR solution structure of rat a*β*(1-16): toward understanding the mechanism of rats’ resistance to Alzheimer’s disease. Biophys J, 102, 136–143. 10.1016/j.bpj.2011.11.4006

56. Brünger, A.T., Adams, P.D., Clore, G.M., et al. (1998) Crystallography & NMR system: A new software suite for macromolecular structure determination. Acta Crystallogr D Biol Crystallogr, 54, 905–921. 10.1107/S0907444998003254

57. Kuszewski, J., Gronenborn, A.M. and Clore, G.M. (1997) Improvements and extensions in the conformational database potential for the refinement of NMR and X-ray structures of proteins and nucleic acids. Journal of Magnetic Resonance, 125, 171–177. 10.1006/jmre.1997.1116

58. Laskowski, R.A., Rullmannn, J.A., MacArthur, M.W., et al. (1996) AQUA and PROCHECK-NMR: programs for checking the quality of protein structures solved by NMR. J Biomol NMR, 8, 477–486. 10.1007/BF00228148

59. Ivanova, E.V., Kolosov, P.M., Birdsall, B., et al. (2007) Eukaryotic class 1 translation termination factor eRF1 − the NMR structure and dynamics of the middle domain involved in triggering ribosome-dependent peptidyl-tRNA hydrolysis. The FEBS Journal, 274, 4223–4237. 10.1111/j.1742-4658.2007.05949.x

60. Goddard, T.D., Huang, C.C., Meng, E.C., et al. (2018) UCSF ChimeraX: Meeting modern challenges in visualization and analysis. Protein Science, 27, 14–25. 10.1002/pro.3235

61. Farrow, N.A., Zhang, O., Forman-Kay, J.D., et al. (1994) A heteronuclear correlation experiment for simultaneous determination of ^15^N longitudinal decay and chemical exchange rates of systems in slow equilibrium. J Biomol NMR, 4, 727–734. 10.1007/BF00404280

62. Polshakov, V.I., Birdsall, B., Frenkiel, T.A., et al. (1999) Structure and dynamics in solution of the complex of *Lactobacillus casei* dihydrofolate reductase with the new lipophilic antifolate drug trimetrexate. Protein Science, 8, 467–481. 10.1110/ps.8.3.467

63. Dosset, P., Hus, J.C., Blackledge, M., et al. (2000) Efficient analysis of macromolecular rotational diffusion from heteronuclear relaxation data. J Biomol NMR, 16, 23–28. 10.1023/a:1008305808620

64. Kuznetsov, D., Tegenfeldt, F., Manni, M., et al. (2023) OrthoDB v11: annotation of orthologs in the widest sampling of organismal diversity. Nucleic Acids Research, 51, D445–D451. 10.1093/nar/gkac998

65. Huerta-Cepas, J., Szklarczyk, D., Heller, D., et al. (2019) eggNOG 5.0: a hierarchical, functionally and phylogenetically annotated orthology resource based on 5090 organisms and 2502 viruses. Nucleic Acids Research, 47, D309–D314. 10.1093/nar/gky1085

66. Wheeler, D.L., Barrett, T., Benson, D.A., et al. (2007) Database resources of the National Center for Biotechnology Information. Nucleic Acids Research, 35, D5–D12. 10.1093/nar/gkl1031

67. Shen, W., Sipos, B. and Zhao, L. (2024) SeqKit2: A Swiss army knife for sequence and alignment processing. iMeta, 3, e191. 10.1002/imt2.191

68. Edgar, R.C. (2004) MUSCLE: multiple sequence alignment with high accuracy and high throughput. Nucleic Acids Research, 32, 1792–1797. 10.1093/nar/gkh340

69. Waterhouse, A.M., Procter, J.B., Martin, D.M.A., et al. (2009) Jalview Version 2—a multiple sequence alignment editor and analysis workbench. Bioinformatics, 25, 1189–1191. 10.1093/bioinformatics/btp033

70. Potter, S.C., Luciani, A., Eddy, S.R., et al. (2018) HMMER web server: 2018 update. Nucleic Acids Research, 46, W200–W204. 10.1093/nar/gky448

71. Yu, D., Chojnowski, G., Rosenthal, M., et al. (2023) AlphaPulldown—a python package for protein–protein interaction screens using AlphaFold-Multimer. Bioinformatics, 39, btac749. 10.1093/bioinformatics/btac749

72. Capella-Gutiérrez, S., Silla-Martínez, J.M. and Gabaldón, T. (2009) trimAl: a tool for automated alignment trimming in large-scale phylogenetic analyses. Bioinformatics, 25, 1972–1973. 10.1093/bioinformatics/btp348

73. Chamberlain,S.A. and Szöcs,E. (2013) taxize: taxonomic search and retrieval in R. F1000Res, 2, 191. 10.12688/f1000research.2-191.v2

74. Minh, B.Q., Schmidt, H.A., Chernomor, O., et al. (2020) IQ-TREE 2: New Models and Efficient Methods for Phylogenetic Inference in the Genomic Era. Molecular Biology and Evolution, 37, 1530–1534. 10.1093/molbev/msaa015

75. Müller, T. and Vingron, M. (2000) Modeling amino acid replacement. J Comput Biol, 7, 761–776. 10.1089/10665270050514918

76. Yang, Z. (1994) Maximum likelihood phylogenetic estimation from DNA sequences with variable rates over sites: approximate methods. J Mol Evol, 39, 306– 314. 10.1007/BF00160154

77. The modENCODE Consortium, Roy, S., Ernst, J., et al. (2010) Identification of functional elements and regulatory circuits by *Drosophila* modENCODE. Science, 330, 1787–1797. 10.1126/science.1198374

78. Öztürk-Çolak, A., Marygold, S.J., Antonazzo, G., et al. (2024) FlyBase: updates to the *Drosophila* genes and genomes database. GENETICS, 227, iyad211. 10.1093/genetics/iyad211

79. Jumper, J., Evans, R., Pritzel, A., et al. (2021) Highly accurate protein structure prediction with AlphaFold. Nature, 596, 583–589. 10.1038/s41586-021-03819-2

80. Blum, M., Andreeva, A., Florentino, L.C., et al. (2025) InterPro: the protein sequence classification resource in 2025. Nucleic Acids Research, 53, D444–D456. 10.1093/nar/gkae1082

81. Cutler, G., Perry, K.M. and Tjian, R. (1998) Adf-1 is a nonmodular transcription factor that contains a TAF-binding Myb-like motif. Molecular and Cellular Biology, 18, 2252–2261. 10.1128/MCB.18.4.2252

82. Wiegmann, B.M., Trautwein, M.D., Winkler, I.S., et al. (2011) Episodic radiations in the fly tree of life. Proc Natl Acad Sci U S A, 108, 5690–5695. 10.1073/pnas.1012675108

83. The UniProt Consortium, Bateman, A., Martin, M.-J., et al. (2025) UniProt: the Universal Protein Knowledgebase in 2025. Nucleic Acids Research, 53, D609–D617. 10.1093/nar/gkae1010

84. Larkin, M.A., Blackshields, G., Brown, N.P., et al. (2007) Clustal W and Clustal X version 2.0. Bioinformatics, 23, 2947–2948. 10.1093/bioinformatics/btm404

85. Filippova, G.N., Fagerlie, S., Klenova, E.M., et al. (1996) An exceptionally conserved transcriptional repressor, CTCF, employs different combinations of zinc fingers to bind diverged promoter sequences of avian and mammalian c-*myc* oncogenes. Molecular and Cellular Biology, 16, 2802–2813. 10.1128/MCB.16.6.2802

86. Laskowski, R.A., Jabłońska, J., Pravda, L., et al. (2018) PDBsum: Structural summaries of PDB entries. Protein Science, 27, 129–134. 10.1002/pro.3289

87. Evans, R., O’Neill, M., Pritzel, A., et al. (2021) Protein complex prediction with AlphaFold-Multimer. 10.1101/2021.10.04.463034. 10.1101/2021.10.04.463034

88. Pavletich, N.P. and Pabo, C.O. (1993) Crystal structure of a five-finger GLI-DNA complex: new perspectives on zinc fingers. Science, 261, 1701–1707. 10.1126/science.8378770

89. Aravind, L. (2000) The BED finger, a novel DNA-binding domain in chromatin-boundary-element-binding proteins and transposases. Trends in Biochemical Sciences, 25, 421–423. 10.1016/S0968-0004(00)01620-0

90. Lannes, L., Furman, C.M., Hickman, A.B., et al. (2023) Zinc-finger BED domains drive the formation of the active *Hermes* transpososome by asymmetric DNA binding. Nat Commun, 14, 4470. 10.1038/s41467-023-40210-3

91. Westman, B.J., Perdomo, J., Matthews, J.M., et al. (2004) Structural studies on a protein-binding zinc-finger domain of Eos reveal both similarities and differences to classical zinc fingers. Biochemistry, 43, 13318–13327. 10.1021/bi049506a

92. Mena, E.L., Kjolby, R.A.S., Saxton, R.A., et al. (2018) Dimerization quality control ensures neuronal development and survival. Science, 362, eaap8236. 10.1126/science.aap8236

93. Bertolini, M., Fenzl, K., Kats, I., et al. (2021) Interactions between nascent proteins translated by adjacent ribosomes drive homomer assembly. Science, 371, 57–64. 10.1126/science.abc7151

94. Nikolova, E.N., Stanfield, R.L., Dyson, H.J., et al. (2020) A Conformational Switch in the Zinc Finger Protein Kaiso Mediates Differential Readout of Specific and Methylated DNA Sequences. Biochemistry, 59, 1909–1926. 10.1021/acs.biochem.0c00253

95. Kabsch, W. and Sander, C. (1983) Dictionary of protein secondary structure: Pattern recognition of hydrogen-bonded and geometrical features. Biopolymers, 22, 2577–2637. 10.1002/bip.360221211

