## Supplementary Information for "Beyond DNA Binding: single C2H2 zinc fingers with adjacent β-strands mediate dimerization in *Drosophila* transcription factors"

**Supplementary Figure S1. NOE distribution. (A)** NOE histograms giving the number of long-range (green, i-j > 4), medium-range (red, 1 < i-j < 4), sequential (blue, i-j = 1) and intra-residue (grey) NOEs for each protein residue in IMZF$^{1\text{-}62}$ structure calculation. **(B)** NOE map of the IMZF$^{1\text{-}62}$. The dots below the diagonal correspond to the NOEs between the protein backbone atoms (Hα and HN). The dots above the diagonal correspond to the observed NOEs between all protons. The figure was generated using the program NMRest, written in-house.

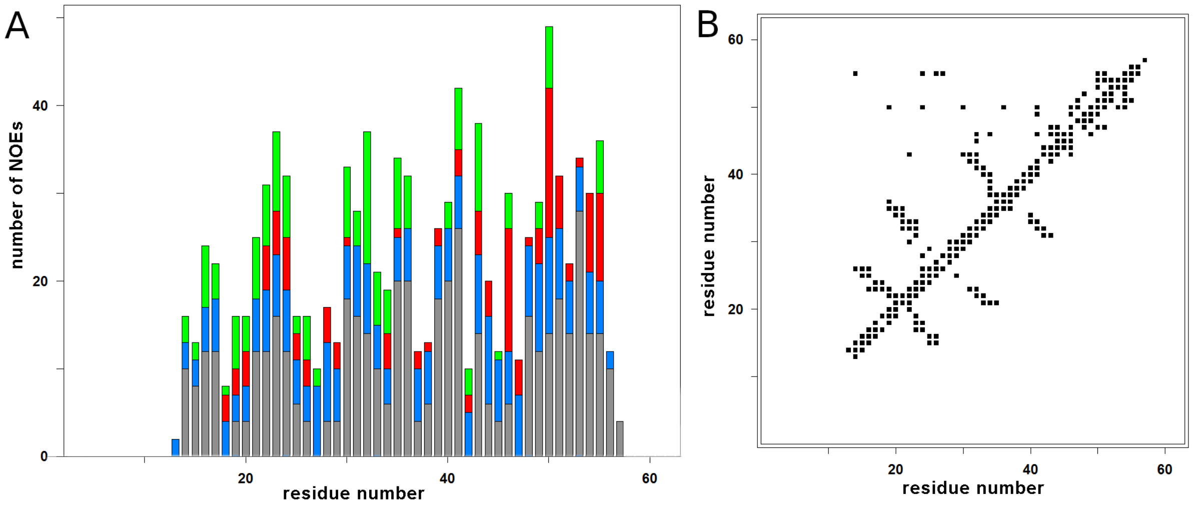

**Supplementary Figure S2. The Ramachandran map plot (φ and ψ torsion angles for the protein backbone).** (**A**) Best conformer (#1) of solution structures of the IMZF$^{1\text{-}62}$ dimer. (**B**) 20 conformers of the NMR families of solution structures of the IMZF$^{1\text{-}62}$ dimer. Glycine residues are marked by triangles and all the other residues are shown as squares.

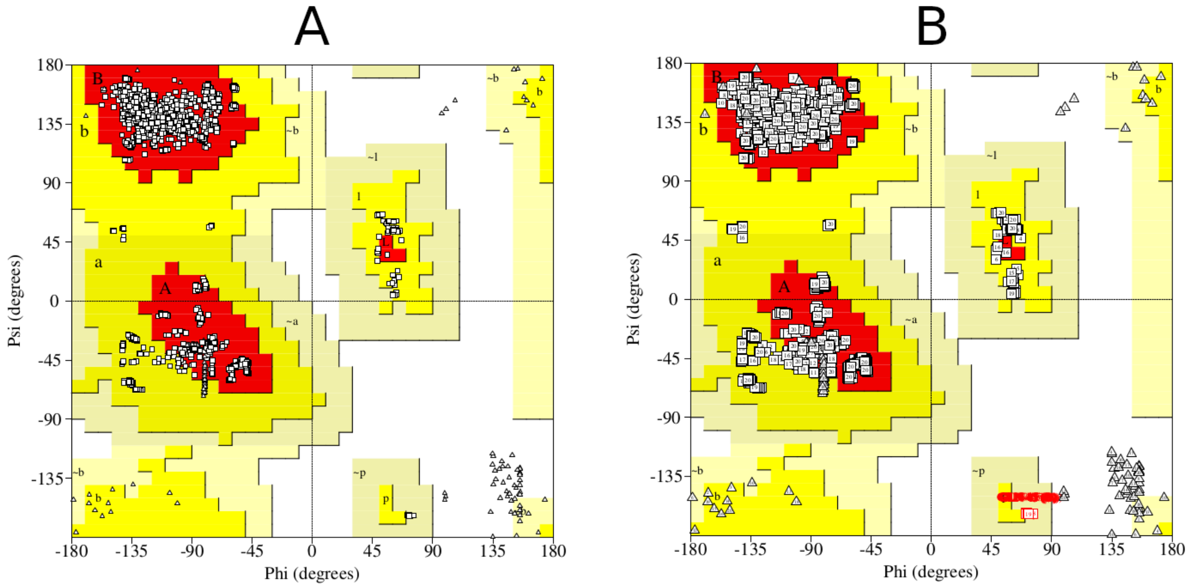

**Supplementary Figure S3. Fragment of a multiple sequence alignment of orthologs containing 'CGEx'-type C2H2 domains.** Amino acid residues are colored with the ClustalX color scheme. Secondary structure elements are labeled according to Stogios et al., 2005. Color key is shown in **Supplementary Table S1**.

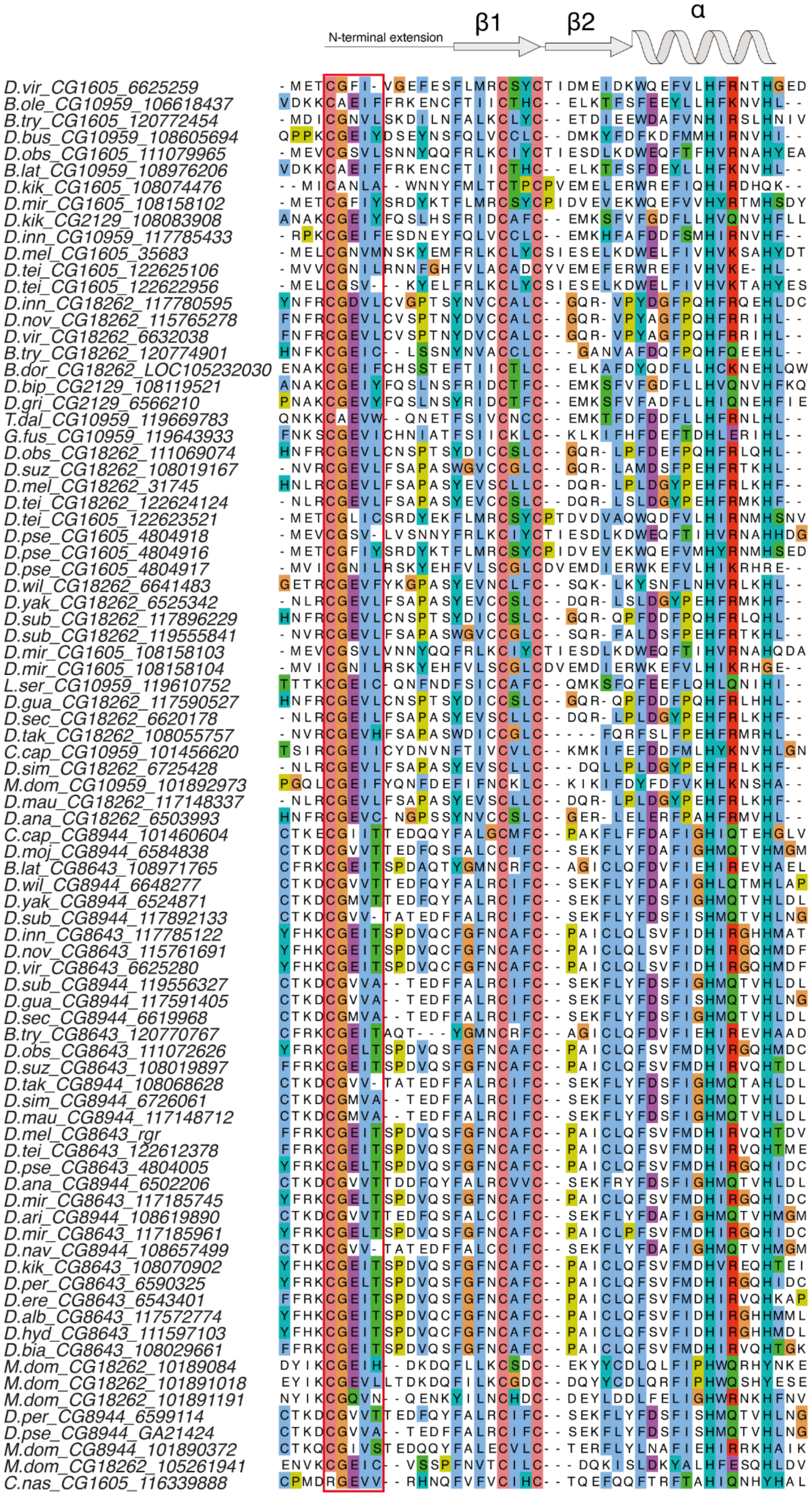

**Supplementary Figure S4.** Phylogenetic reconstruction of orthologous proteins containing ‘CGEx’-type C2H2 domains.

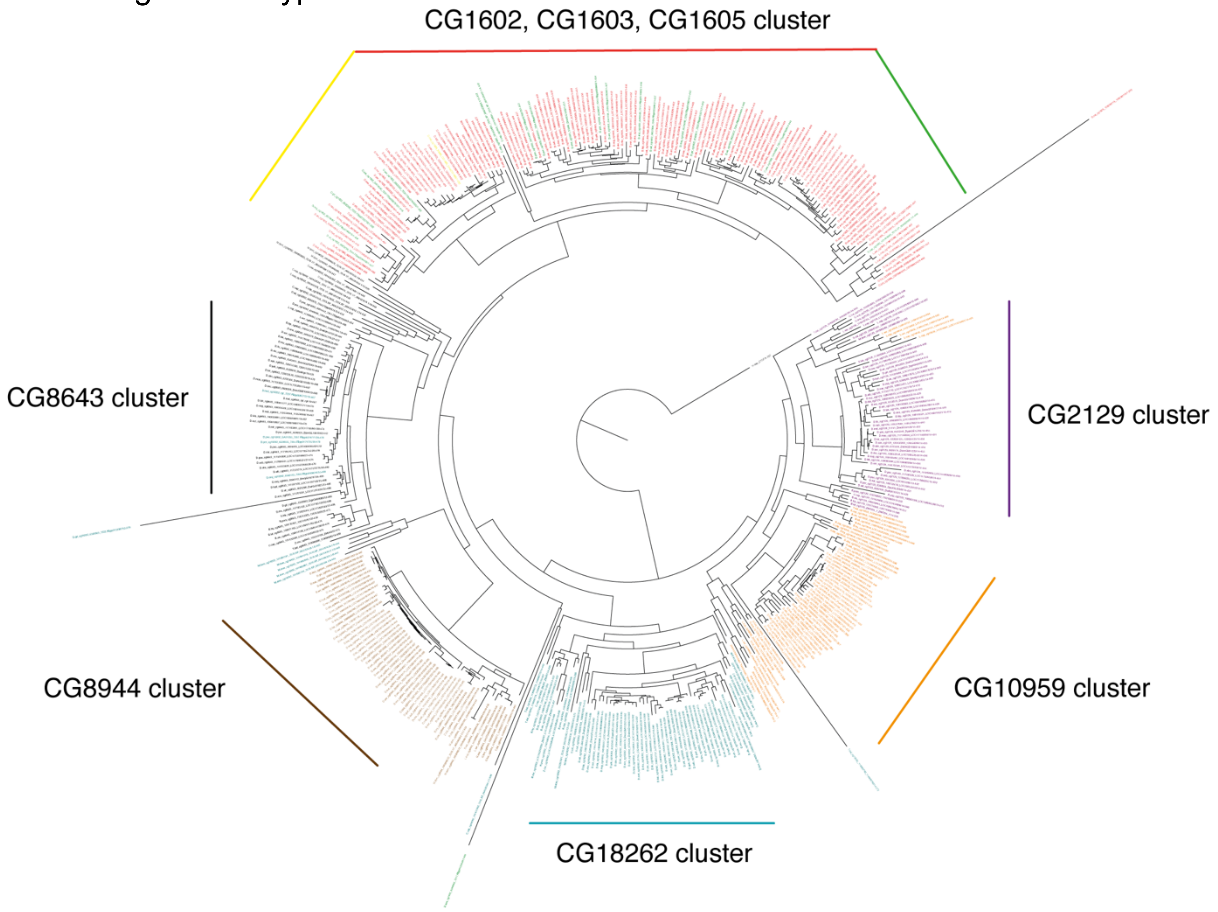

**Supplementary Figure S5. Results of yeast two-hybrid assays.** Growth assay plates without histidine are shown (yeasts are unable to grow on this medium in the absence of interaction). AD stands for Activation Domain, BD for DNA-Binding Domain of GAL4 protein.

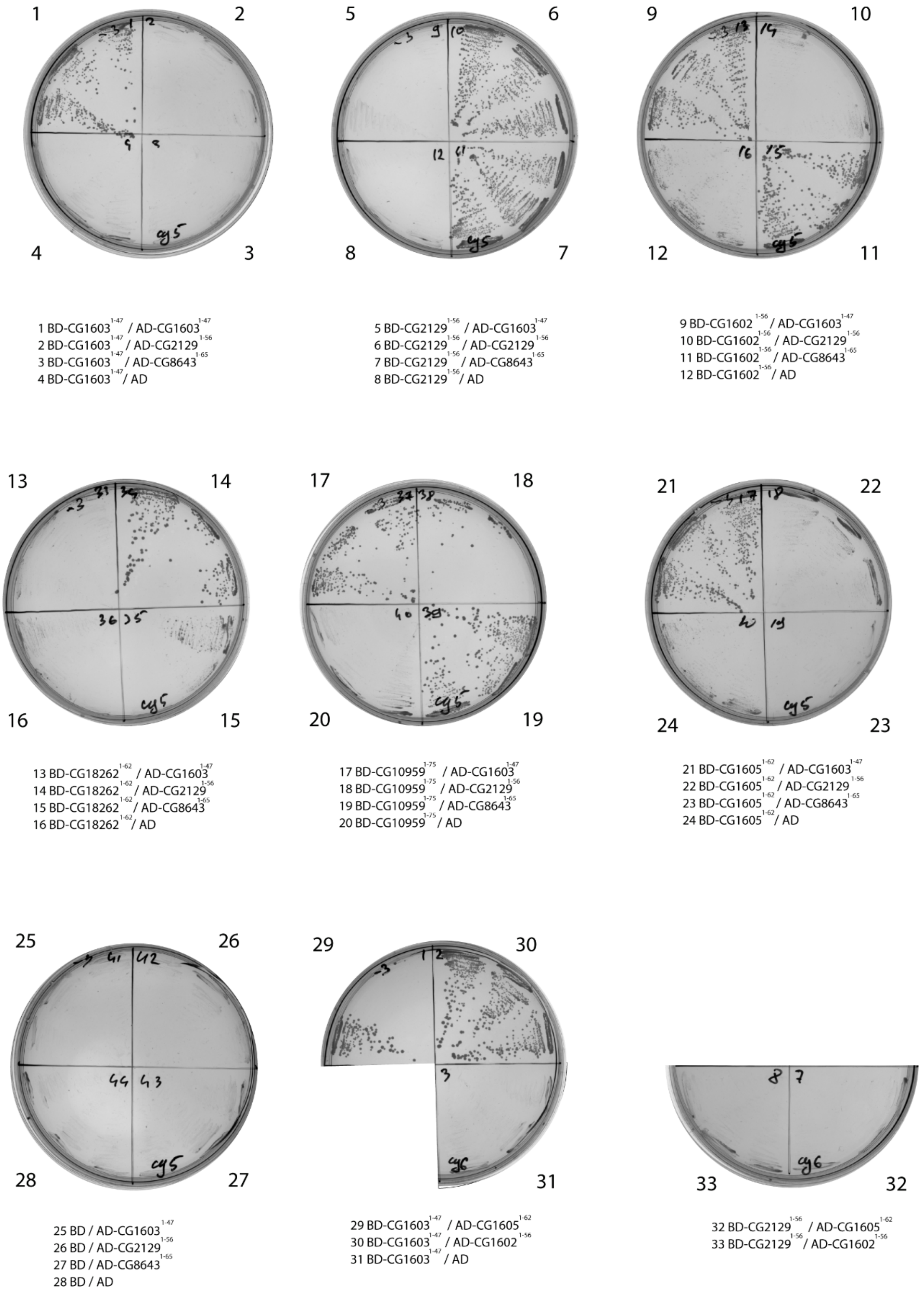

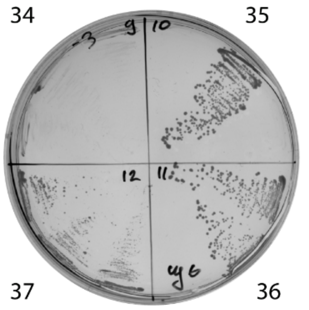

34 BD-CG2129$^{1-56}$ / AD
35 BD-CG1602$^{1-56}$ / AD-CG1605$^{1-62}$
36 BD-CG1602$^{1-56}$ / AD-CG1602$^{1-56}$
37 BD-CG1602$^{1-56}$ / AD

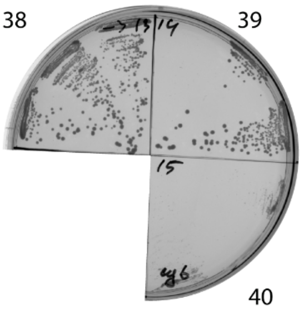

38 BD-CG1605$^{1-62}$ / AD-CG1605$^{1-62}$
39 BD-CG1605$^{1-62}$ / AD-CG1602$^{1-56}$
40 BD-CG1605$^{1-62}$ / AD

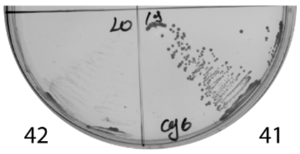

41 BD-CG18262$^{1-62}$ / AD-CG1605$^{1-62}$
42 BD-CG18262$^{1-62}$ / AD-CG1602$^{1-56}$

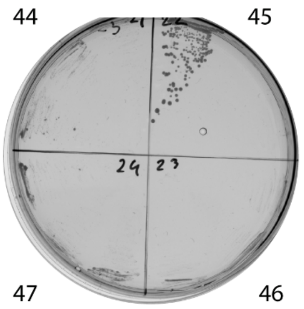

44 BD-CG18262$^{1-62}$ / AD
45 BD-CG10959$^{1-75}$ / AD-CG1605$^{1-62}$
46 BD-CG10959$^{1-75}$ / AD-CG1602$^{1-56}$
47 BD-CG10959$^{1-75}$ / AD

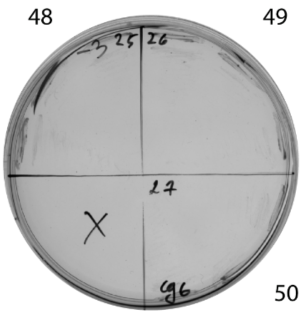

48 BD / AD-CG1605$^{1-62}$
49 BD / AD-CG1602$^{1-56}$
50 BD / AD

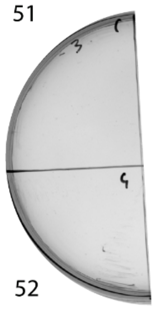

51 BD-CG1603$^{1-47}$ / AD-CG8944$^{1-80}$
52 BD-CG1603$^{1-47}$ / AD

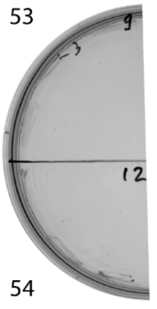

53 BD-CG2129$^{1-56}$ / AD-CG8944$^{1-80}$
54 BD-CG2129$^{1-56}$ / AD

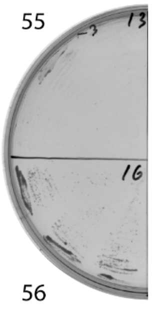

55 BD-CG1602$^{1-56}$ / AD-CG8944$^{1-80}$
56 BD-CG1602$^{1-56}$ / AD

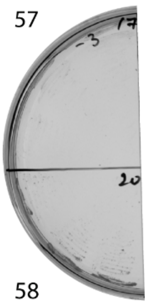

57 BD-CG1605$^{1-62}$ / AD-CG8944$^{1-80}$
58 BD-CG1605$^{1-62}$ / AD

59

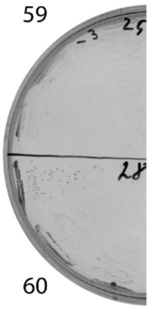

60

59 BD-CG18262$^{1-62}$ / AD-CG8944$^{1-80}$
60 BD-CG18262$^{1-62}$ / AD

61

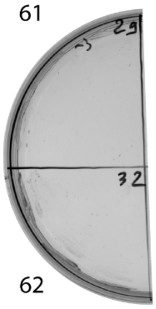

62

61 BD-CG10959$^{1-75}$ / AD-CG8944$^{1-80}$
62 BD-CG10959$^{1-75}$ / AD

63

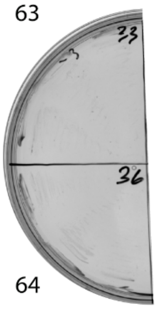

64

63 BD / AD-CG8944$^{1-80}$
64 BD / AD

65

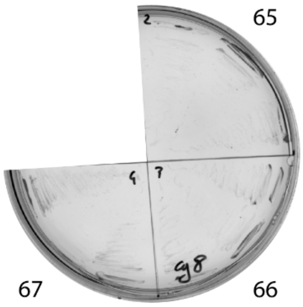

67 66

65 BD-CG1603$^{1-47}$ / AD-CG18262$^{1-62}$
66 BD-CG1603$^{1-47}$ / AD-CG10959$^{1-75}$
67 BD-CG1603$^{1-47}$ / AD

68

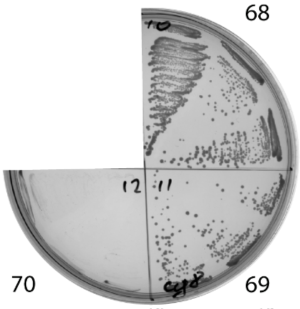

70 69

68 BD-CG2129$^{1-56}$ / AD-CG18262$^{1-62}$
69 BD-CG2129$^{1-56}$ / AD-CG10959$^{1-75}$
70 BD-CG2129$^{1-56}$ / AD

71

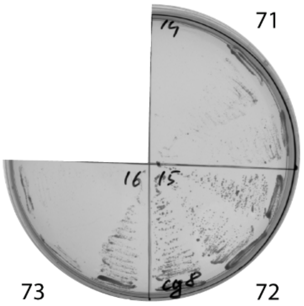

73 72

71 BD-CG1602$^{1-56}$ / AD-CG18262$^{1-62}$
72 BD-CG1602$^{1-56}$ / AD-CG10959$^{1-75}$
73 BD-CG1602$^{1-56}$ / AD

74

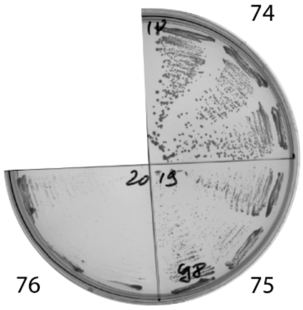

76 75

74 BD-CG1605$^{1-62}$ / AD-CG18262$^{1-62}$
75 BD-CG1605$^{1-62}$ / AD-CG10959$^{1-75}$
76 BD-CG1605$^{1-62}$ / AD

77

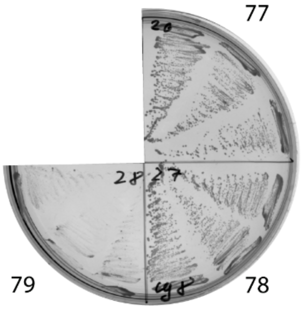

79 78

77 BD-CG18262$^{1-62}$ / AD-CG18262$^{1-62}$
78 BD-CG18262$^{1-62}$ / AD-CG10959$^{1-75}$
79 BD-CG18262$^{1-62}$ / AD

80

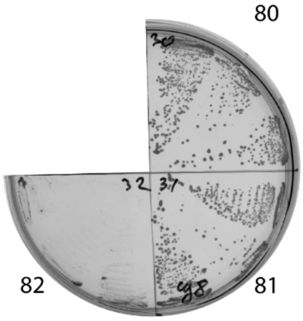

82 81

80 BD-CG10959$^{1-75}$ / AD-CG18262$^{1-62}$
81 BD-CG10959$^{1-75}$ / AD-CG10959$^{1-75}$
82 BD-CG10959$^{1-75}$ / AD

83

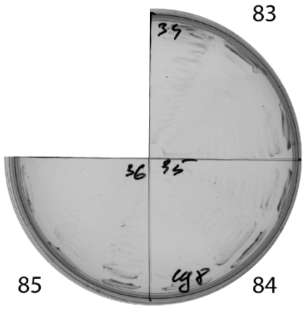

85 84

83 BD / AD-CG18262$^{1-62}$
84 BD / AD-CG10959$^{1-75}$
85 BD / AD

**Supplementary Figure S6. Hydrogen/deuterium exchange protection in the IMZF$^{1\text{-}62}$ dimer.** The representative NMR structure is colored according to H/D exchange rates: green indicates residues with slow exchange (protection for >24 h), blue denotes intermediate exchange (1–24 h), and uncolored regions (if applicable) reflect faster exchange.

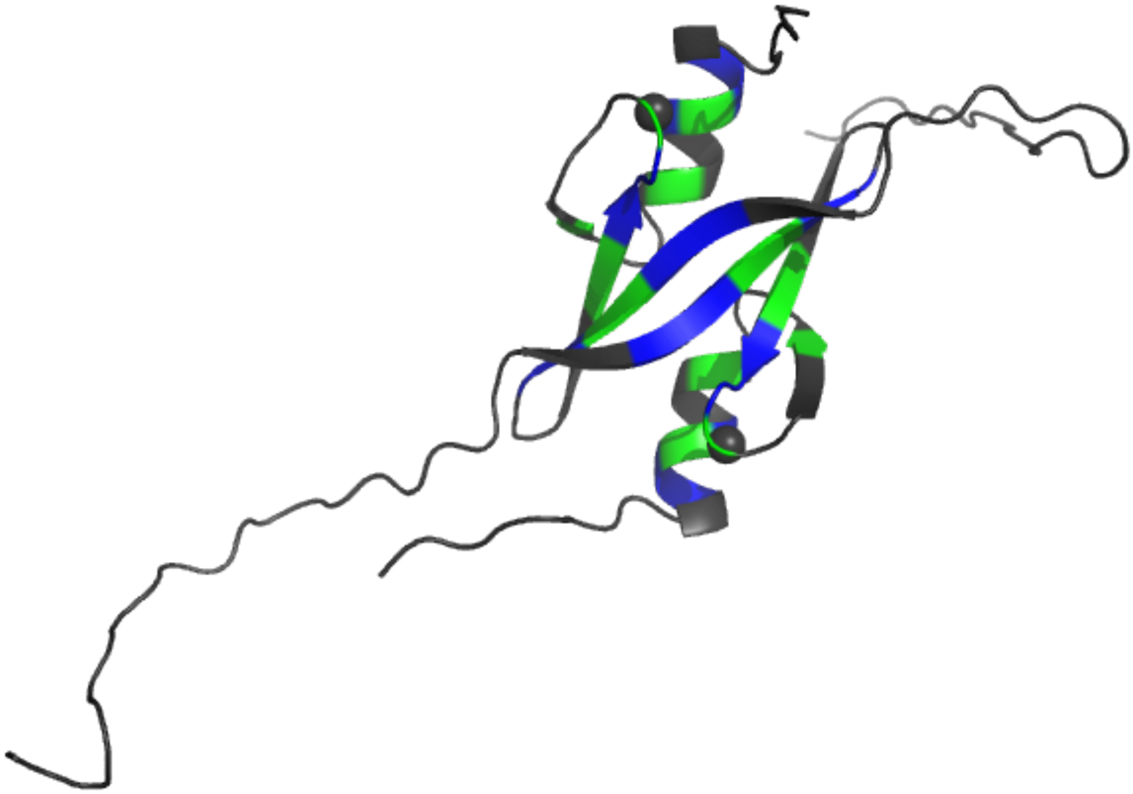

**Supplementary Figure S7. The relaxation parameters of the amide $^{15}N$ nuclei of each residue of the IMZF$^{1\text{-}62}$ dimer, measured at 600 MHz proton resonance frequency and 298 K.** (**A**) The heteronuclear $^{15}N,^{1}H$-steady-state NOE values. (**B**) The longitudinal relaxation rate $R_1$ ($s^{-1}$). (**C**) The transverse relaxation rate $R_2$ ($s^{-1}$). (**D**) The order parameter $S^2$ determined by model-free analysis. (**E**) Chemical exchange $R_{ex}$ contributions to the transverse relaxation rates ($s^{-1}$).

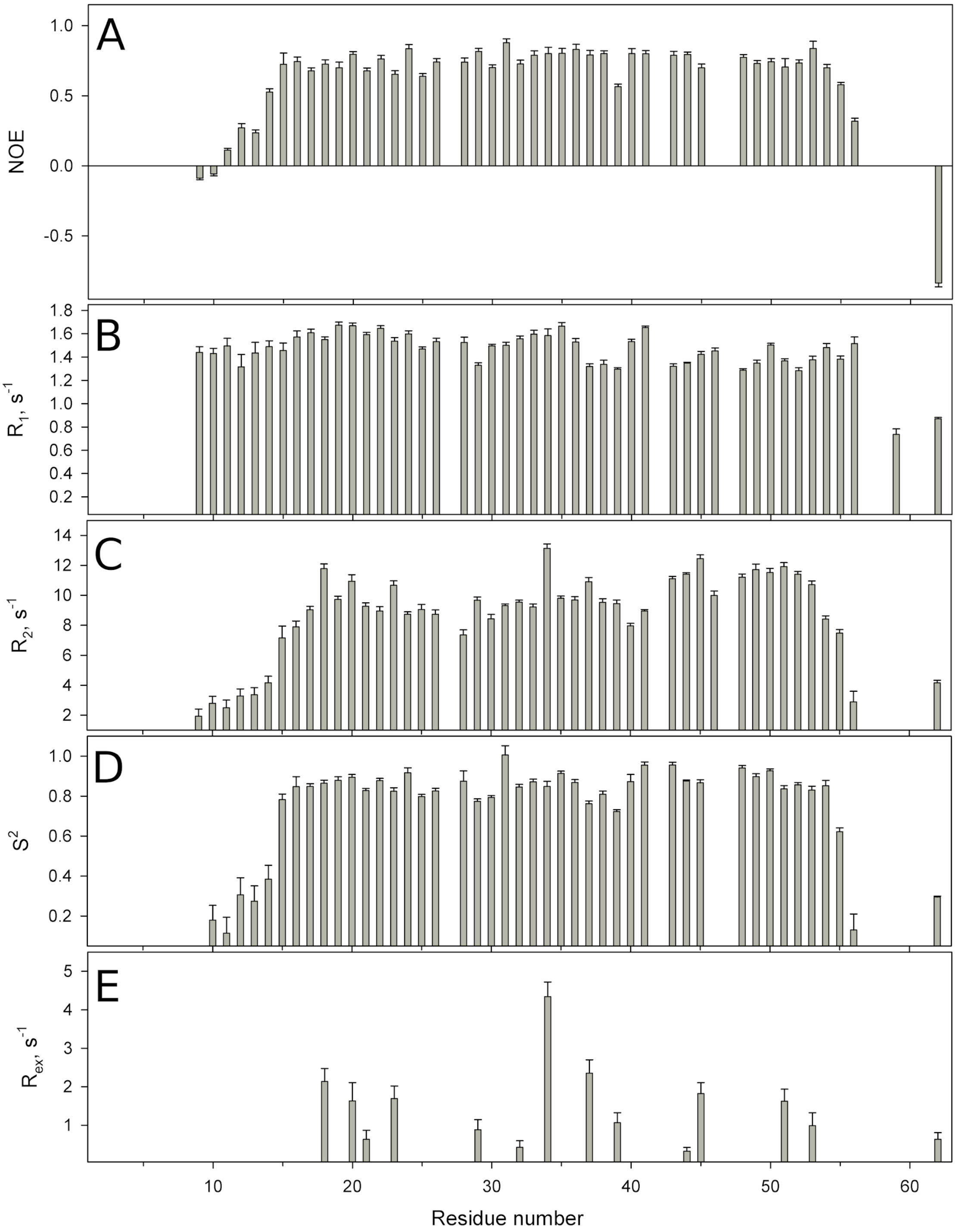

**Supplementary Figure S8. Representative NMR structure of IMZF$^{1\text{-}62}$ dimer is shown according to protein mobility in a broad time scale.** The thickness of the chain reflects the value ($1 - S^2$) representing the extent of the local amplitude of protein backbone motion in ps-ns time scale. Color-coded kinetics: orange – residues undergoing conformational exchange in ms time scale (with $R_{ex} > 2\ s^{-1}$); blue – slow H/D exchange (1h); green – very slow H/D exchange (24h).

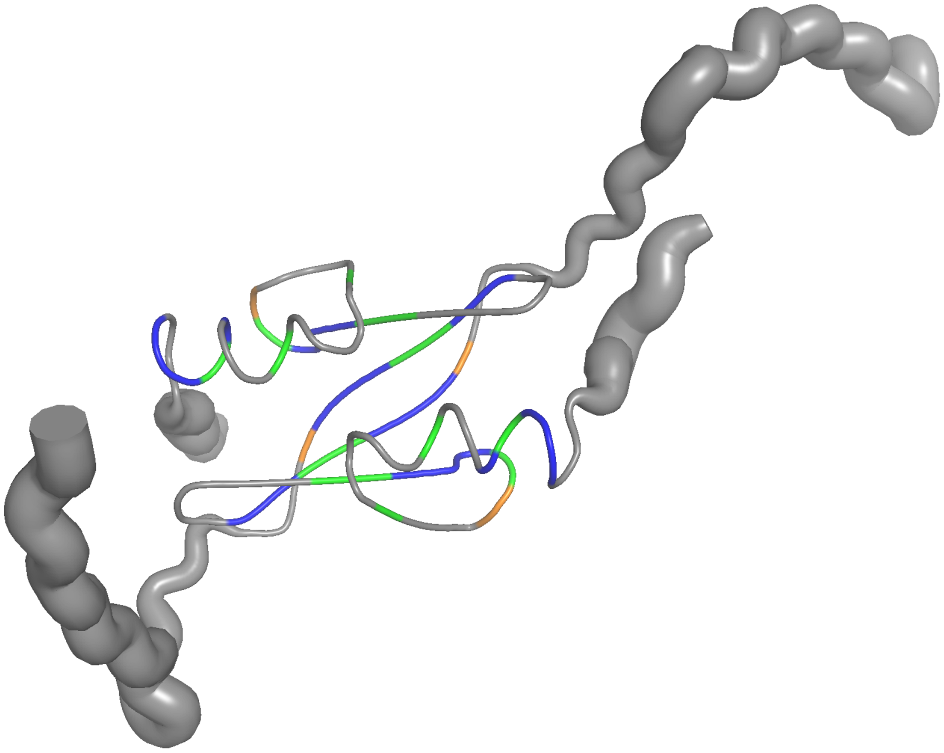

**Supplementary Figure S9. Anisotropic rotational diffusion of the IMZF$^{1-62}$ dimer.** The ellipsoid represents the dimer's rotational diffusion anisotropy, with its axes scaled according to the experimentally derived diffusion tensor ($D_a/D_x \approx 2.0$). One subunit is color-coded based on the order parameters of residues with experimentally measured NMR data.

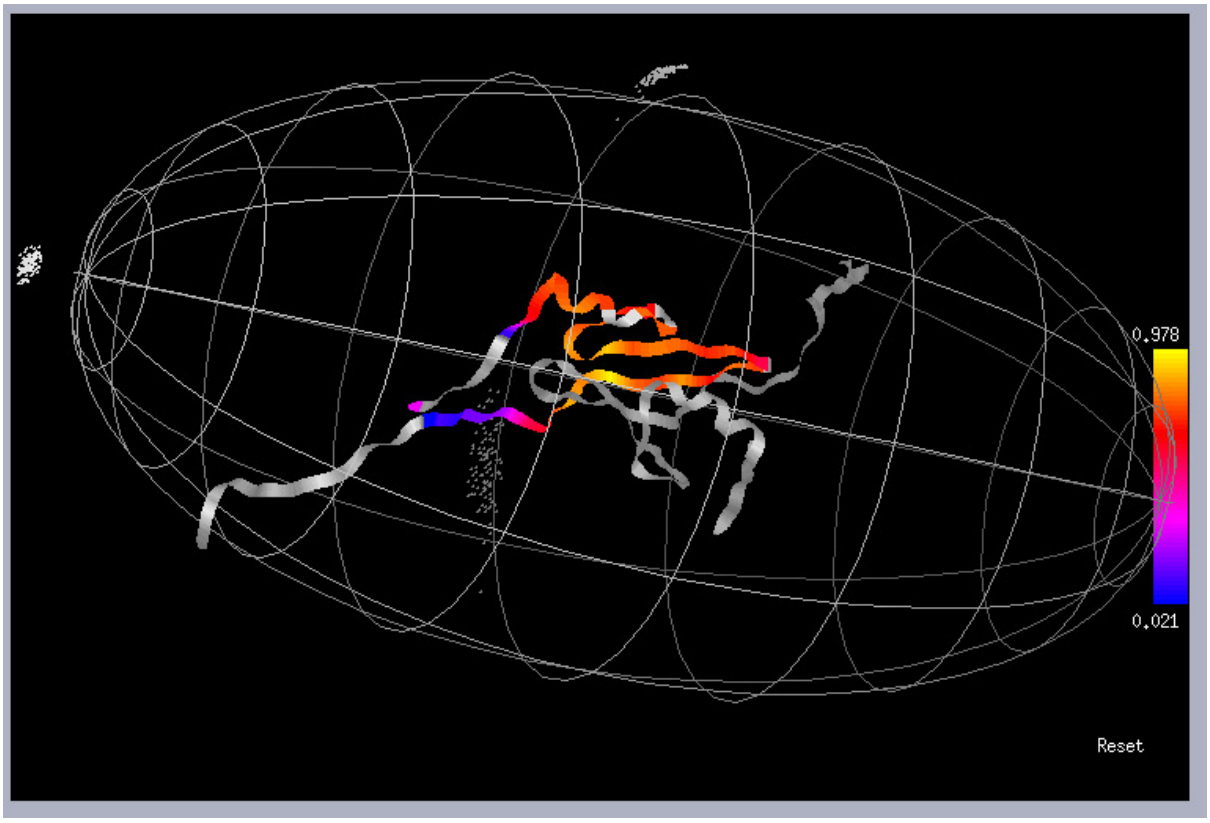

**Supplementary Figure S10. Alphafold2 models of all CGEx-type C2H2 domains from *D. melanogaster*.** The top row of each pair of models is colored according to confidence prediction by pLDDT (the confidence scale is displayed to the right of the models), while the bottom row is colored by chain. Input sequences for the models were: CG1602$^{1\text{-}56}$, Mzfp1$^{1\text{-}47}$, CG1605$^{1\text{-}62}$, CG2129$^{1\text{-}56}$, CG10959$^{1\text{-}75}$, IMZF$^{1\text{-}62}$, CG8643$^{1\text{-}65}$, and CG8944$^{1\text{-}80}$. The amino acid ranges displayed in the figure are: CG1602$^{1\text{-}43}$, Mzfp1$^{1\text{-}43}$, CG1605$^{1\text{-}44}$, CG2129$^{14\text{-}56}$, CG10959$^{21\text{-}61}$, IMZF$^{16\text{-}54}$, CG8643$^{20\text{-}58}$, and CG8944$^{18\text{-}59}$.

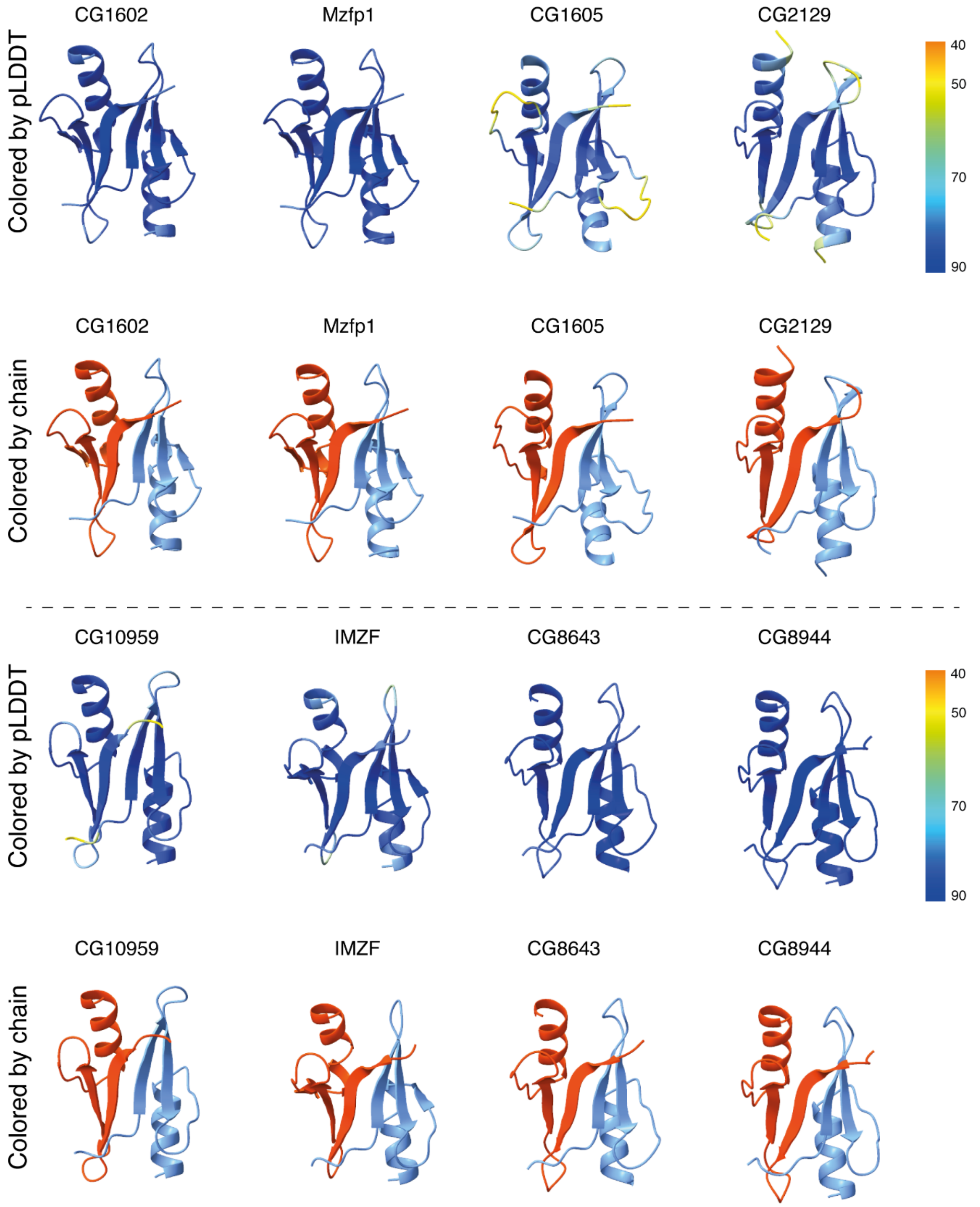

**Supplementary Table S1**. **Oligonucleotides used for cloning.** Restriction enzyme sites are shown in small letters; the corresponding enzymes are noted. Nucleotide substitutions in mutagenic primers are also shown in small letters.

| Name | Sequence | Restriction enzyme |
|---|---|---|
| 1603d | AACgaattcATGGAGCTGTGCGGC | *EcoRI* |
| 1603_47r | TTTAgtcgacGTCTTCGCAGTAGTGCG | *SalI* |
| 2129d | GCCgaattcATGCTCAAGTCCCTCAAG | *EcoRI* |
| 2129_56r | TTTAgtcgacCTCGAAGTGAATGTTCTGG | *SalI* |
| 1602d | TAAgaattcATGGAAACTTGCGGCC | *EcoRI* |
| 1602_56r | TTAgtcgacCACTGGCTCGAAGTCG | *SalI* |
| 1605d | AAGgaattcATGGTGGTTTGCGGG | *EcoRI* |
| 1605_62r | TTAgtcgacAATGACTTCGGAAGTAGTGAG | *SalI* |
| 8944_d | ACTggatccATGAGTGTCCTGGCGTATCC | *BamH* |
| 8944_80r | TTCctcgagTTACATGGGCTCGCCGAGATTG | *XhoI* |
| CG18262_d | AACgaattCATGATCAAGGTGCAGCCACC | *EcoRI* |
| 18262_62r | TTTAgtcgacCAGCGAGCTGGAGCTGTTC | *SalI* |
| CG10959_d | AACgaattcATGTTGAGACACTGCCTCGC | *EcoRI* |
| 10959_75r | TTTActcgagTCCCGTGACCGTCTTGGTGA | *XhoI* |
| CG8643_d | AACgaattcATGATGCCGCTGCAGGAGTC | *EcoRI* |
| 8643_65r | TTTAgtcgacGCTCACGTCCTGCGTGTGCT | *SalI* |
| 8643_65rS | TTTAgtcgacTTAGCTCACGTCCTGCGTGTGCT | *SalI* |
| 18262_62rS | TTTAgtcgacTTACAGCGAGCTGGAGCTGTTC | *SalI* |
| 10959_75rS | TTTActcgagTTATCCCGTGACCGTCTTGGTGA | *XhoI* |
| 18262_26d | AACggatccGCACCCGCCTCCTACGA | *BamH* |
| 18262Y30Ed | GCCTCCgaaGAAGTGAGCT | |
| 18262Y30Er | CTCACTTCttcGGAGGCGGG | |

| | |
|---|---|
| 18262F50Ad | GGAACACgccCGGCTGAAACAC |
| 18262F50Ar | TCAGCCGggcGTGTTCCGGGTA |
| 18262F50Ed | GGAACACgaaCGGCTGAAACAC |
| 18262F50Er | TCAGCCGttcGTGTTCCGGGTA |

**Supplementary Table S2. Statistics for the ensemble of the calculated 20 structures of the IMZF$^{1\text{-}62}$.**

(**A**). The number and distribution of restraints. The total number of restraints used for the two polypeptide chains is indicated.

| **Total NOEs** | 1674 | **Total H-bonds** | 34 |
|---|---|---|---|
| Long range (\|i − j\|> 4) | 162 | Intrachain | 28 |
| Medium (1 < \|i − j\|≤ 4) | 132 | Interchain | 6 |
| Sequential (\|i − j\|= 1) | 268 | **Dihedral angles** | 278 |
| Intraresidue | 1034 | **Chemical Shifts** | 84 |
| Interchain | 78 | **Zn-cooordinations** | 8 |

(**B**). Restraint violations and structural statistics (for 20 structures).

| **Average RMSD** | **Ensemble of 20 final structures** | **Representative structure** |
|---|---|---|
| **From experimental restraints** | | |
| Distance (Å) | 2.27 ± 0.06 | 2.32 |
| Dihedral (°) | 41.8 ± 0.1 | 41.8 |
| **From idealized covalent geometry** | | |
| Bonds (Å) | 0.0126 ± 0.0001 | 0.0127 |
| Angles (°) | 0.95 ± 0.02 | 0.95 |
| Impropers (°) | 0.82 ± 0.02 | 0.82 |
| **Ramachandran plot statistics** | | |
| % of residues in most favorable region of Ramachandran plot | 82.4 | 81.2 |
| % of residues in disallowed region of Ramachandran plot | 0 | 0 |

| | |
|---|---|
| (**C**). Superimposition on the representative structure (Å). | |
| Backbone (C, Cα, N) RMSD over the structured protein core (residues 13-55) | 0.52 ± 0.25 |

**Table S3.** Distribution of orthologs from C2H2 transcription factor family containing 'CGEx'-type C2H2 domains.

| Family | | Number of species | Order |
|---|---|---|---|
| **1** | Cecidomyiidae | 1 | Diptera |
| **2** | Glossinidae | 3 | Diptera |
| **3** | Calliphoridae | 2 | Diptera |
| **4** | Muscidae | 2 | Diptera |
| **5** | Tephritidae | 8 | Diptera |
| **6** | Drosophilidae | 38 | Diptera |
| **7** | Diopsidae | 1 | Diptera |

**Supplementary Table S4.** Clustal X Colour Scheme.

| **Clustal X Default Colouring** | | | |
|---|---|---|---|
| **Category** | **Colour** | **Residue at position** | **{ Threshold, Residue group }** |
| Hydrophobic | BLUE | A,C,I,L,M,F,W,V | {>60%, WLVIMAFCYHP} |
| | | C | {>60%, WLVIMAFCYHP} |
| Positive charge | RED | K,R | {>60%,KR},{>85%, K,R,Q} |
| Negative charge | MAGENTA | E | {>60%,KR},{>50%,QE},{>50%,ED},{>85%,E,Q,D} |
| | | D | {>60%,KR}, {>85%, D,E,N}, {>50%,ED} |
| Polar | GREEN | N | {>50%, N}, {>85%, N,D} |
| | | Q | {>60%,KR},{>50%,QE},{>85%,Q,T,K,R} |
| | | S,T | {>60%, WLVIMAFCYHP}, {>50%, TS}, {>85%,S,T} |
| Cysteines | PINK | C | {>85%, C} |
| Glycines | ORANGE | G | {>0%, G} |
| Prolines | YELLOW | P | {>0%, P} |
| Aromatic | CYAN | H,Y | {>60%, WLVIMAFCYHP}, {>85%, W,Y,A,C,P,Q,F,H,I,L,M,V} |
| Unconserved | WHITE | any / gap | If none of the above criteria are met |

**Table S5. The list of fragments of C2H2 domain proteins that were cloned to compatible vectors for Y2H.** The same fragments were cloned into modified Pet32a vector (Novagen) for bacterial expression.

| Protein | UniProt ID | Sequence | Y2H vector |
|---|---|---|---|
| CG18262$^{1-62}$ | Q9W3J0 | MIKVQPPPDVAGGYHNLRCGEVLFSAPASYEVSCLLCDQRLPLDGYPEHFRLKHFTNSSSSL | pGAD, pGBT |
| CG10959$^{1-75}$ | Q9W3J2 | MLRHCLANVDYVRLHAQQQLRPKCGEIFYEPEANRFQLVCLLCDMKHFGFEDFARHIRNVHFDKQGRPLTKTVTG | pGAD, pGBT |
| CG8643$^{1-65}$ | E1JH10 | MMPLQESPSPSWQLEDYAFFRKCGEITVSPDVQSFGFNCAFCPAICLQFSVFMDHIRVQHTQDVS | pGAD, pGBT |
| CG1603$^{1-47}$ | Q95TL6 | MELCGNVMTNSKYEMFRLKCLYCSIESELKDWELFIVHVKSAHYCED | pGAD, pGBT-C |
| CG1602$^{1-56}$ | Q8MSB3 | METCGLICVSRDYEKFLMRCSYCPTDVEVAQWQEFVLHIRNVHSKVRSMEDDFEPV | pGAD, pGBT |
| CG1605$^{1-62}$ | Q5BIC3 | MVVCGNILARNNFEHFVLACAHQNDCYVEMEFKRWREFIVHVKEHLEANPSMQQELTTSEVI | pGAD, pGBT |
| CG2129$^{1-56}$ | Q9W3J4 | MLKSLKPDTYAGMANAKCGEIYFQSLHSFRIDCAFCEMKSFVFGDFLLHVQNIHFE | pGAD, pGBT-C |
| CG8944$^{1-80}$ | Q9VXQ6 | MSVLAYPRTNDEEIFRHCTKDCGMVTATDDFQYFALRCIFCSEKFLYFDSFIGHMQTVHLGDQSVANTSSRLPFNLGEPM | pGAD, pGBT |

**Supplementary Table S6.** The full list of mutations in the C2H2 domain of IMZF protein tested by SEC-MALS.

| Protein | Mutation | Impact on oligomeric state (SEC-MALS) |
|---|---|---|
| cg18262$^{26\text{-}62}$ | 26-62 | + |
| cg18262$^{1\text{-}62}$ | Y30E | + |
| cg18262$^{1\text{-}62}$ | F50A | - |
| cg18262$^{1\text{-}62}$ | F50E | + |

**Supplementary Table S7. The presence of C2H2 domains with additional N-terminal secondary structure elements across proteomes of higher Eukaryota.** The confidence of dimers predicted by AlphaFold (AF) were assessed based on inter-chain PAE score < 5.0 and pLDDT score over 70 for interacting domains.

| | Additional β-strand | | | Additional α-helix | Total ZFs |
|---|---|---|---|---|---|
| Species | Total | Dimers predicted by AF | Dimers like IMZF | Total | |
| *Acyrthosiphon pisum* | 33 | 16 | 2 | 267 | 4843 |
| *Amphimedon queenslandica* | 8 | 3 | 0 | 27 | 342 |
| *Arabidopsis thaliana* | 9 | 3 | 0 | 65 | 439 |
| *Bombyx mori* | 124 | 70 | 0 | 170 | 4572 |
| *Caenorhabditis elegans* | 31 | 10 | 0 | 117 | 892 |
| *Capitella teleta* | 24 | 9 | 0 | 223 | 2770 |
| *Ciona intestinalis* | 9 | 2 | 0 | 40 | 1308 |
| *Crassostrea gigas* | 20 | 9 | 0 | 125 | 2815 |
| *Danio rerio* | 35 | 15 | 0 | 171 | 21268 |
| *Daphnia pulex* | 21 | 9 | 0 | 77 | 1058 |
| *Dictyostelium discoideum* | 5 | 2 | 0 | 29 | 104 |
| *Drosophila melanogaster* | 35 | 11 | 6 | 161 | 2325 |
| *Gallus gallus* | 17 | 10 | 0 | 69 | 3997 |
| *Glycine max* | 30 | 11 | 0 | 137 | 771 |
| *Helobdella robusta* | 13 | 5 | 0 | 110 | 1256 |
| *Homo sapiens* | 51 | 22 | 0 | 322 | 12711 |
| *Hydra vulgaris* | 3 | 0 | 0 | 24 | 953 |
| *Ixodes scapularis* | 11 | 4 | 0 | 98 | 1594 |
| *Lingula anatina* | 29 | 10 | 0 | 130 | 3690 |
| *Monosiga brevicollis* | 10 | 4 | 0 | 34 | 204 |
| *Mus musculus* | 38 | 10 | 0 | 261 | 10496 |
| *Octopus bimaculoides* | 48 | 14 | 1 | 428 | 22528 |
| *Oryza sativa* | 11 | 3 | 0 | 55 | 566 |
| *Strigamia maritima* | 12 | 8 | 0 | 70 | 1370 |
| *Trichoplax adhaerens* | 3 | 1 | 0 | 23 | 217 |
| *Xenopus laevis* | 56 | 29 | 0 | 231 | 10212 |
| *Zea mays* | 11 | 2 | 0 | 36 | 703 |
